# Protein Phosphatase 2A Activation Attenuates Acute Myocardial Injury in Takotsubo Syndrome by Modulating Ferroptosis and Mitochondrial Injury in Cardiomyocytes

**DOI:** 10.64898/2026.07.27.740349

**Authors:** Ti Wang, Qian Xu, Jisheng Sun, Hildebrando Candido Ferreira Neto, Feng Dong, Colin Stomberski, Caitlin M. O’Connor, Goutham Narla, Daxin Wang, Zhiyong Lin

## Abstract

**Background:** Takotsubo syndrome (TTS) is an acute stress-induced cardiomyopathy characterized by transient left ventricular dysfunction. Despite its reversible nature, TTS is associated with substantial morbidity and mortality in the acute phase, and no specific treatments are available. The precise molecular mechanisms that connect catecholamine stress to reversible myocardial injury are not fully understood. PP2A, a holoenzyme with serine/threonine phosphatase activity, plays a vital role in normal cardiac development, and its dysregulation has been associated with heart disease. Its role in TTS and its potential as a therapeutic target remain completely unknown.

**Methods:** Analysis of public multi-omics datasets from stress cardiomyopathy (SCM) and experimental models of TTS, along with treatment of cardiomyocytes with human TTS plasma, was used to investigate a potential role for protein phosphatase 2A (PP2A) in stress-induced myocardial injury. To clarify the functional impact of manipulating PP2A activity in TTS, we used a series of disease relevant cell based and in vivo models, leveraging both genetic and pharmacological approaches to modulate PP2A activity in cardiomyocytes and in mice. To gain mechanistic insights into how PP2A influences TTS pathology and downstream signaling pathways, RNA sequencing, stress-responsive iron handling, mitochondrial function, and cardiac phenotypes were thoroughly evaluated in both in vivo and in vitro studies.

**Results:** PP2A activity was markedly reduced in cardiac tissues from mice with isoprenaline-induced TTS, as well as in isoprenaline-treated cardiomyocytes. Genetic or pharmacological inhibition of PP2A worsened catecholamine-induced cardiac dysfunction and myocardial injury. Most notably, pharmacological activation of PP2A using an orally bioavailable small-molecule activator strongly mitigated myocardial damage and enhanced cardiac function in TTS models. Mechanistically, PP2A inactivation promoted JNK-MAPK signaling and dysregulated stress-responsive iron-handling pathways, leading to ferritinophagy-mediated ferroptosis and mitochondrial dysfunction. Pharmacological JNK inhibition effectively rescued myocardial injury caused by PP2A deficiency in two TTS animal models and in cardiomyocytes.

**Conclusions:** PP2A inactivation is a key molecular event linking catecholamine stress to myocardial injury in TTS. Restoring PP2A activity or inhibiting downstream JNK attenuates ferritinophagy-dependent stress responses and mitochondrial dysfunction, providing a unifying mechanistic framework and highlighting the PP2A-JNK axis as a potential target for short-term intervention during the acute phase of TTS.

**CLINICAL PERSPECTIVE:** *What Is New?:* • Protein phosphatase 2A (PP2A) activity is acutely suppressed during the early phase of Takotsubo syndrome (TTS), with preferential involvement of the apical myocardium, closely mirroring the disease’s characteristic clinical phenotype. • Downstream of PP2A inactivation, catecholamine stress triggers JNK-MAPK signaling activation and dysregulated stress-responsive iron handling, including ferritinophagy-mediated ferroptosis and mitochondrial dysfunction in cardiomyocytes. • Pharmacological restoration of PP2A activity or downstream inhibition of JNK mitigates acute myocardial injury and preserves cardiac function, thereby establishing the PP2A-JNK axis as a central stress-responsive pathway in TTS.

*What Are the Clinical Implications?:* • Identification of PP2A inactivation as a key molecular event highlights a novel opportunity for targeted intervention during the acute phase. • Studies in preclinical animal TTS models provide the first evidence that illuminates the therapeutic potential of a small-molecule PP2A activator as a strategy for acute TTS and may serve as a broad cardioprotective strategy in settings where cardiac dysfunction is implicated. • Targeting downstream JNK signaling, which is activated by PP2A inactivation, emerges as an additional mechanism-based strategy for attenuating acute myocardial injury in TTS.

## INTRODUCTION

Takotsubo syndrome (TTS) is an acute stress-induced cardiomyopathy characterized by transient left ventricular dysfunction that often mimics acute coronary syndrome.^1–3^ Although once considered a benign condition, accumulating evidence indicates that TTS is associated with substantial morbidity and mortality during the acute phase, including heart failure, arrhythmias, and cardiogenic shock.^4,5^ Although TTS and acute coronary syndromes have similar in-hospital mortality rates, they are distinct conditions. TTS differs from acute coronary syndromes in that it does not involve obstructed coronary arteries. Additionally, TTS is characterized by a unique pattern of anteroseptal-apical dyskinetic ballooning. Despite its prevalence, no disease-specific therapies currently exist, and clinical management remains largely supportive.^6,7^ The molecular mechanisms by which acute stress induces myocardial injury in TTS, while allowing subsequent functional recovery, remain poorly understood, and therapies targeting the underlying drivers of disease pathogenesis are lacking.

Excessive catecholamine exposure is widely recognized as a central trigger of TTS, yet the downstream molecular events linking catecholamine stress to cardiomyocyte injury remain poorly defined.^7–9^ Prior studies have implicated β-adrenergic signaling, calcium overload, oxidative stress, microvascular dysfunction, and metabolic disturbances in the pathophysiology of TTS.^10–14^ More recently, stress-responsive kinase signaling pathways have been suggested to contribute to acute myocardial injury.^15^ However, far less is known about whether corresponding alterations in phosphatase activity contribute to TTS pathogenesis, particularly during the early and reversible phase of the disease. Protein phosphatase 2A (PP2A) is a major serine/threonine phosphatase that regulates multiple stress-response pathways, including MAPK signaling, metabolic homeostasis, and mitochondrial function.^16–18^ Dysregulation of PP2A activity has been implicated in a range of cardiovascular diseases, including heart failure, ischemia-reperfusion injury, and atherosclerosis, underscoring its importance in maintaining myocardial signaling homeostasis.^19–22^ More importantly, pharmacological tools that selectively modulate PP2A activity, such as small-molecule PP2A activators, have been reported to confer protection in several cardiovascular disease models.^22–25^ However, whether PP2A activity is dynamically regulated during TTS and how such regulation influences cardiomyocyte vulnerability remains unexplored.

In this study, we identify PP2A inactivation as a central molecular event in acute TTS. By integrating analyses of multi-omics datasets, patient plasma, animal models, and mechanistic studies in cardiomyocytes, we demonstrate that acute catecholamine stress induces rapid, region-specific suppression of PP2A activity. Loss of PP2A function during the acute phase promotes activation of JNK-MAPK signaling and coordinated dysregulation of stress-responsive iron handling, accompanied by ferroptosis-mediated ferritinophagy, mitochondrial dysfunction, and cardiomyocyte injury. Importantly, pharmacological restoration of PP2A activity or downstream inhibition of JNK effectively mitigates myocardial injury across different experimental systems. Together, these findings establish PP2A as a critical integrator of stress-responsive signaling in the heart and provide a unifying mechanistic framework for the reversible myocardial injury characteristic of TTS.

## MATERIALS AND METHODS

Detailed experimental procedures and additional methodological information are provided in the Supplemental Material.

### Human plasma samples

Human plasma samples from patients with TTS were obtained at Emory University Hospital under protocols approved by the Institutional Review Board of Emory University. Plasma samples from individuals without known cardiovascular or other diseases served as controls. A total of 5 control and 5 TTS plasma samples were included in this study. Blood samples were collected in EDTA-containing tubes, centrifuged to isolate plasma, aliquoted, and stored at -80°C until use.

### Animals

All protocols involving animals were approved by an Institutional Animal Review Committee at Emory University and were performed in compliance with the Guide for the Care and Use of Laboratory Animals. Male C57BL/6J mice aged 16 weeks were purchased from the Jackson Laboratory (Bar Harbor, ME). Cardiac-specific PP2A-Cα deficient mice were generated by crossing the floxed Ppp2cα mice (purchased from Nanjing Biomedical Research Institute) with Myh6-Cre^+/-^ mice (The Jackson Laboratory, strain no. 005657). Cardiac-specific Ppp2cα deficiency was achieved by administering 2 consecutive doses of tamoxifen (50μl, 20 mg/mL, dissolved in sunflower seed oil; Sigma-Aldrich) through intraperitoneal injection to 14-week-old male Ppp2cα flox^+/-^-Myh6-Cre^+/-^ mice (CM-PP2Afl^+/-^). Age-matched male Ppp2cα-floxed mice without Cre expression and Myh6-Cre^+/-^ mice, subjected to the same tamoxifen regimen, were used as controls. Following a 2-week washout period, 16-week-old CM-PP2Afl^+/-^ mice were used for the modeling of TTS.

### Statistical Analysis

Statistical analyses were performed using GraphPad Prism version 9.0 (GraphPad Software Inc, San Diego, CA, USA). The data from the quantitative experiments are presented as mean ± standard error of the mean (SEM) from at least 3 independent experiments. For comparisons between 2 groups, an unpaired 2-tailed Student’s t test was performed for normally distributed data, and the Mann-Whitney test for non-normally distributed data. For comparisons among multiple groups, one-way ANOVA (followed by Tukey post hoc test) was applied. The survival rates were assessed using Kaplan-Meier analysis. P < 0.05 was considered statistically significant.

## RESULTS

### Conserved dysregulation of phosphorylation-dependent signaling in TTS suggests impaired PP2A activity

To elucidate the molecular mechanisms underlying acute cardiac injury in TTS, we first conducted integrative analyses of publicly available transcriptomic and proteomic datasets across multiple experimental models and human samples. Gene Ontology (GO) enrichment analysis of a publicly available rat stress cardiomyopathy (SCM) bulk RNA-seq dataset (GSE223385)^26^ revealed prominent enrichment of stress-responsive biological processes, including extracellular matrix organization, stress-activated MAPK signaling, inflammatory responses, and mitochondrial-related pathways (Figure 1A). Consistently, our RNA-seq analysis of cardiac tissues from a mouse isoprenaline-induced TTS model revealed significant enrichment of pathways associated with programmed cell death, MAPK cascade activation, protein phosphorylation, and inflammatory responses (Figure 1B). To determine whether these signaling features were specific to catecholamine excess or reflected a broader stress cardiomyopathy phenotype, we analyzed data from transverse aortic constriction (TAC)-induced TTS in potassium channel Kv1.5-/- mice.^27^ Notably, GO enrichment analysis again highlighted regulation of protein phosphorylation and activation of MAPK cascades (Figure 1C), suggesting that stress-responsive kinase signaling represents a conserved molecular feature across distinct models of stress-induced cardiac dysfunction. More importantly, analysis of a human proteomic dataset from female patients with stress cardiomyopathy (GSE95368)^28^ demonstrated similar enrichment of stress-related signaling pathways, including MAPK family signaling, post-translational protein phosphorylation, programmed cell death, and mitochondrial biogenesis (Figure 1D). Together, these cross-species and cross-model analyses indicate that TTS is characterized by conserved dysregulation of phosphorylation-dependent stress signaling.

**Figure 1.**
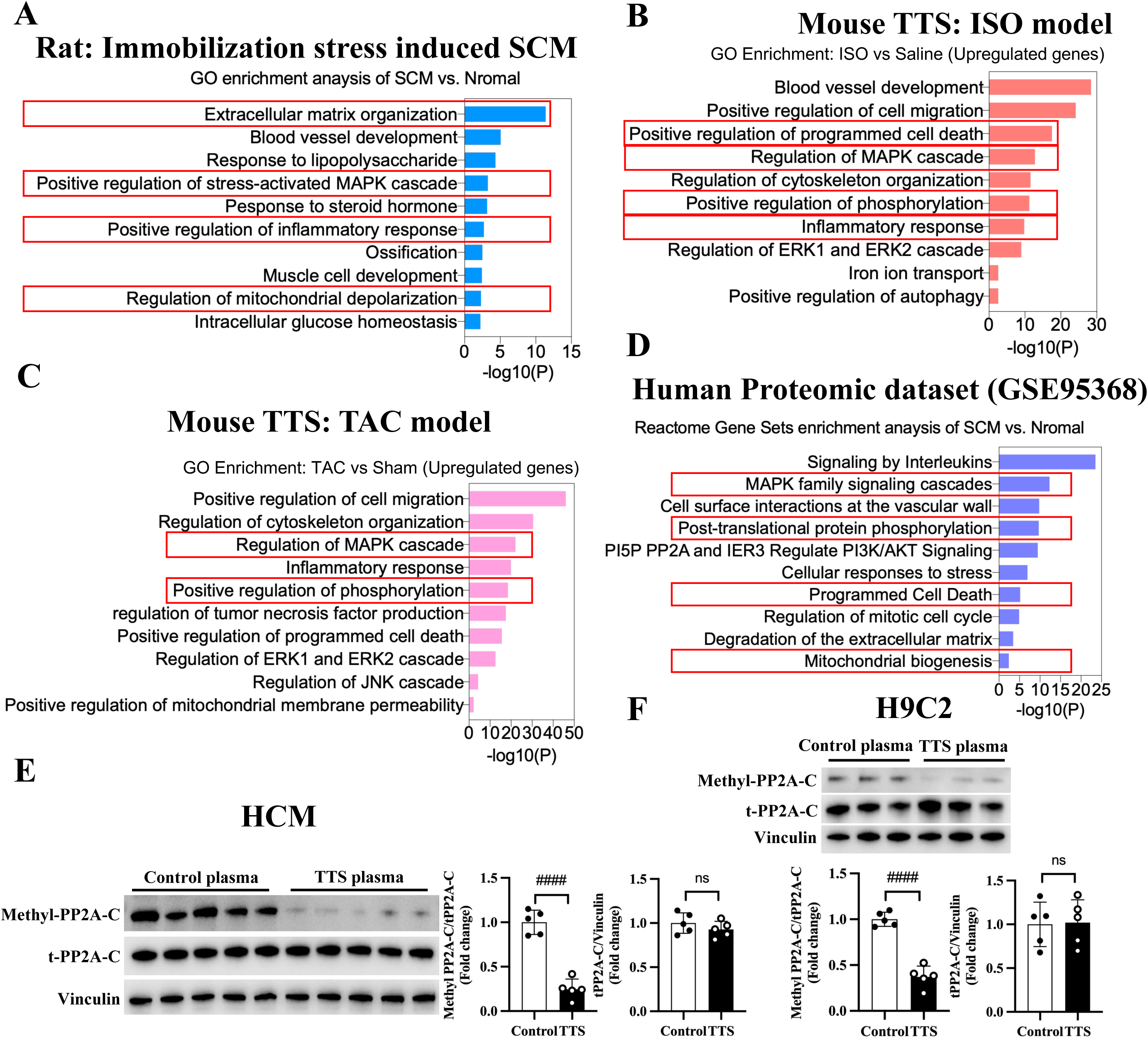
Conserved dysregulation of phosphorylation signaling associates with reduced PP2A activity in TTS. A, Gene Ontology (GO) enrichment analysis of differentially expressed genes in rat bulk RNA sequencing data from an immobilization stress-induced stress cardiomyopathy (SCM) model compared with control rats. **B,** GO enrichment analysis of upregulated genes in a mouse isoproterenol (ISO)-induced Takotsubo syndrome (TTS) model compared with saline-treated controls. **C,** GO enrichment analysis of upregulated genes in a mouse transverse aortic constriction (TAC) model compared with sham-operated controls. **D,** Reactome pathway enrichment analysis of differentially expressed proteins in the human SCM proteomic dataset. **E,** Western blot and quantification data of methylated PP2A catalytic subunit (methyl-PP2A-C) and total PP2A-C levels in primary human cardiomyocytes treated with plasma from control subjects or patients with acute TTS (n=5). **F,** Western blot and quantification data of methyl-PP2A-C and total PP2A-C levels in H9C2 cardiomyocytes treated with plasma from control subjects or patients with TTS (n=5). Data are shown as mean ± SD. SCM indicates stress cardiomyopathy; TTS, Takotsubo syndrome; ISO, isoproterenol; TAC, transverse aortic constriction; PP2A, protein phosphatase 2A. #*P*<0.05; ##*P*<0.01; ###*P*<0.001; ####*P*<0.0001; ns, not significant.

To further characterize cellular heterogeneity and signaling alterations at single-cell resolution, we analyzed single-cell RNA sequencing data (GSE305273)^29^ from control and ISO-treated mouse hearts. Uniform manifold approximation and projection (UMAP) identified major cardiac cell populations, including cardiomyocytes, endothelial cells, fibroblasts, and immune cells (Supplementary Figure 1A). Comparison between control and ISO-treated hearts revealed discernible shifts in the distribution of multiple cell populations following catecholamine stress (Supplementary Figure 1B), indicating that TTS is associated with broad remodeling of the cardiac cellular landscape. In particular, stress-responsive cell populations, including cardiomyocytes, fibroblasts, and endothelial cells, exhibited notable changes in their spatial distribution, suggesting differential sensitivity to catecholamine-induced injury. Cell-type composition analysis further revealed distinct shifts in the relative abundance of major cardiac cell populations following ISO treatment (Supplementary Figure 1C). Pathway enrichment analysis of differentially expressed genes demonstrated that ISO treatment was associated with suppression of protein phosphorylation–related processes and dysregulation of MAPK signaling, along with alterations in cell death and mitochondrial pathways (Supplementary Figure 1D). Notably, gene set enrichment analysis further identified coordinated alterations in phosphatase-related pathways (Supplementary Figure 1E). The PP2A catalytic subunit gene “*Ppp2ca*” was broadly expressed across cardiac cell populations and enriched in cardiomyocytes, where its expression was reduced following ISO treatment (Supplementary Figure 1F). Consistently, suppression of phosphorylation-related pathways together with activation of MAPK signaling and inflammation response support a model in which impaired PP2A activity contributes to stress-induced signaling dysregulation in TTS (Supplementary Figure 1G).

Guided by integrative cross-species bioinformatic analyses implicating PP2A, a major serine/threonine phosphatase that regulates cardiac phosphorylation homeostasis, as a key regulator disrupted in TTS, we next sought to determine whether the disease milieu modulates PP2A activity. Because direct measurement of enzymatic activity in human cardiac tissue is not feasible, we used patient plasma as a surrogate to model the systemic disease environment. Plasma samples from patients with acute TTS were applied to primary human cardiomyocytes and H9C2 cells. Compared with plasma from control subjects, acute exposure to TTS plasma significantly reduced methylation of the PP2A catalytic subunit (methyl-PP2A-C), a modification required for full holoenzyme activity, without altering total PP2A protein levels. These findings indicate that circulating factors in TTS patients acutely suppress PP2A activity in cardiomyocytes, supporting the hypothesis that reduced PP2A activity contributes to the pathophysiology of TTS.

Together, these data identify dysregulated phosphorylation signaling as a conserved feature of Takotsubo syndrome across species and disease models and suggest that an acute reduction in PP2A activity is a candidate upstream event that contributes to stress-induced cardiac dysfunction.

### Acute suppression of PP2A activity in the early phase of ISO-induced TTS

Building on our integrative analyses identifying dysregulated phosphorylation signaling and reduced PP2A activity in TTS, we next investigated the temporal and spatial dynamics of PP2A regulation in an established isoproterenol (ISO)-induced acute TTS mouse model. To define the time course of PP2A regulation during disease onset and recovery, mice were analyzed at 2 hours, 7 days, and 14 days after ISO administration. Notably, intraperitoneal administration of ISO (400 mg/kg) rapidly induced a stress cardiomyopathy phenotype in C57BL/6J mice, with a significant increase in acute mortality early after ISO challenge that progressively declined at later time points, consistent with a transient, self-limited disease course (Figure 2A and Supplementary Figure 2A). Echocardiographic assessment 2 hours after ISO administration revealed typical features of TTS, including apparent apical akinesis and basal hyperkinesis of the left ventricle at end-systole, together with a marked reduction in left ventricular systolic function, as evidenced by decreased ejection fraction (EF) and fractional shortening (FS) (Figure 2B). Moreover, echocardiographic analyses demonstrated progressive recovery of cardiac function over time, with partial improvement at 7 days and near-complete normalization by 14 days after ISO challenge (Supplementary Figure 2B). Consistent with the echocardiographic findings, histological analysis revealed prominent myocardial injury during the acute phase after ISO administration. At 2 hours, hematoxylin and eosin (HE) staining showed marked myocardial damage predominantly localized to the left ventricular apex and mid-ventricle, whereas the basal region was relatively preserved (Figure 2C), closely recapitulating the regional vulnerability observed in human TTS. In contrast, myocardial architecture was substantially restored at 7 days and largely normalized by 14 days after ISO administration, paralleling the recovery of cardiac function (Supplementary Figure 2C).

**Figure 2.**
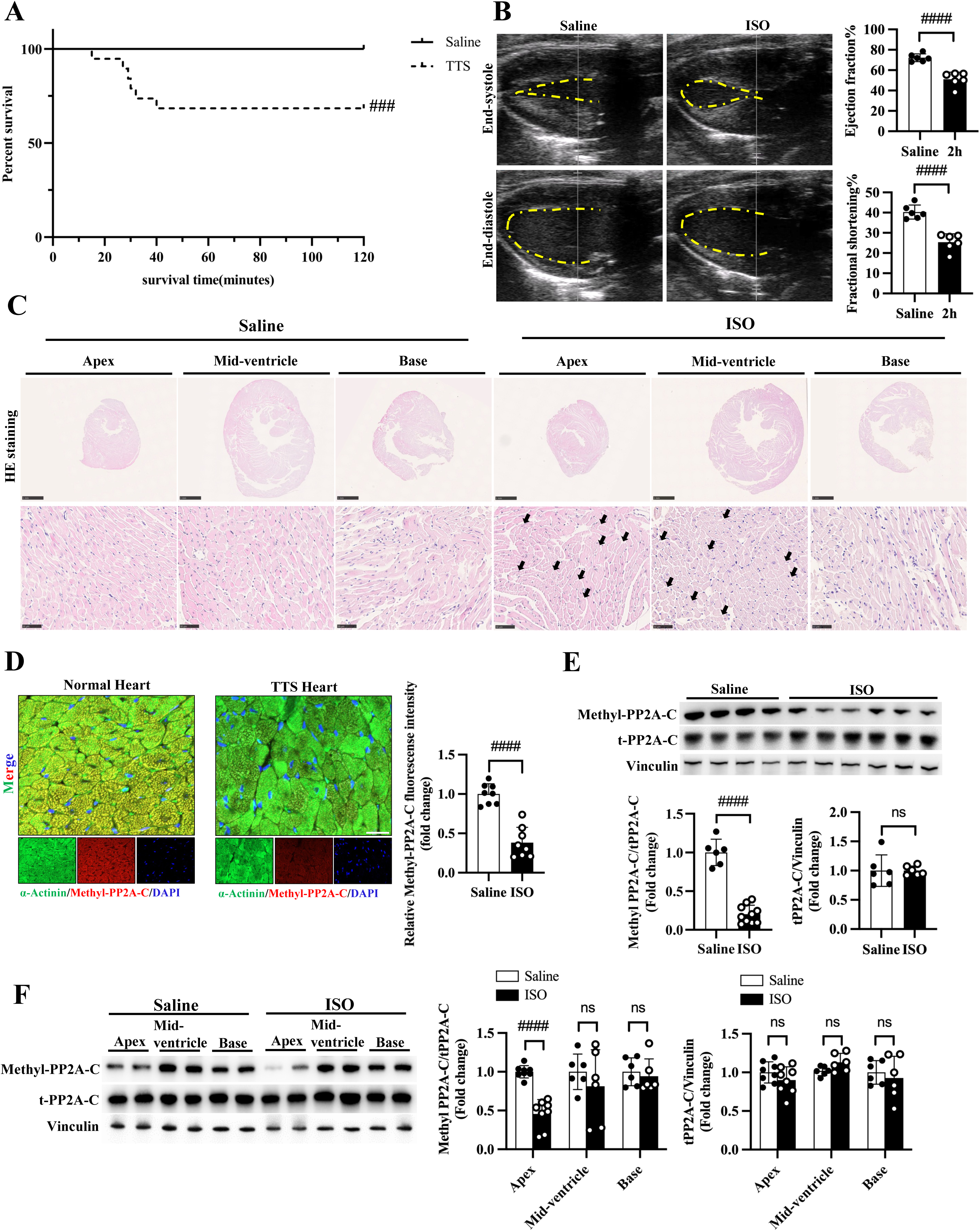
Acute and region-specific suppression of PP2A activity in ISO-induced TTS. A, Kaplan– Meier survival analysis of mice following intraperitoneal administration of ISO. **B,** Representative echocardiographic images at end-systole and end-diastole from saline-and ISO-treated mice at 2 hours, with quantification of left ventricular ejection fraction (EF) and fractional shortening (FS). **C,** Hematoxylin and eosin (HE) staining of left ventricular sections from saline- and ISO-treated mice, showing myocardial injury across apical, mid-ventricular, and basal regions. Arrows indicate areas of myocardial damage. **D,** Representative immunofluorescence staining of methylated PP2A catalytic subunit (methyl-PP2A-C) in cardiomyocytes from saline- and ISO-treated hearts after 2 hours, with quantification of fluorescence intensity. **E,** Immunoblot analysis of methyl-PP2A-C and total PP2A-C (t-PP2A-C) in cardiac tissue from saline- and ISO-treated mice, with quantification normalized to vinculin. **F,** Regional analysis of methyl-PP2A-C and total PP2A-C protein levels in apical, mid-ventricular, and basal myocardium following ISO treatment. Data are presented as mean ± SEM. #*P*<0.05; ##*P*<0.01; ###*P*<0.001; ####*P*<0.0001; ns, not significant.

To determine whether the observed temporal and spatial changes in cardiac function and histology were associated with alterations in PP2A regulation, we next examined PP2A activity at the molecular level. Immunofluorescence staining demonstrated a pronounced reduction in methyl-PP2A-C in cardiomyocytes from ISO-treated hearts at 2 hours, compared with saline-treated controls (Figure 2D). Furthermore, immunoblot analysis revealed that methyl-PP2A-C was markedly suppressed during the acute phase following ISO administration but gradually recovered at 7 and 14 days, in parallel with improvements in cardiac function and resolution of myocardial injury (Figure 2E and Supplementary Figure 2D). In contrast, total PP2A-C protein levels remained stable throughout the disease course, indicating that ISO-induced stress primarily impairs PP2A activity. Given the marked regional heterogeneity of myocardial injury in the ISO-induced TTS model, we next assessed whether PP2A dysregulation exhibited similar spatial specificity. Segmental analysis revealed that reductions in methyl-PP2A-C were most pronounced in the apical myocardium, whereas mid-ventricular and basal regions showed relatively modest changes (Figure 2F). Importantly, total PP2A-C expression remained stable throughout the disease course, further supporting the notion that PP2A activity is transiently and region-specifically suppressed during the acute phase of TTS.

Collectively, these findings demonstrate that acute and region-specific suppression of PP2A activity is a defining molecular feature of ISO-induced TTS, temporally aligned with disease onset and spatially concordant with characteristic apical myocardial dysfunction. This establishes PP2A inactivation as a potential early driver of TTS pathogenesis and provides a mechanistic basis for subsequent functional and molecular investigations.

### Genetic and pharmacological suppression of PP2A activity exacerbates ISO-induced TTS

To determine whether reduced PP2A activity plays a causal role in the development of TTS, we next explored both genetic and pharmacological approaches to suppress PP2A activity in vivo. Cardiomyocyte-specific heterozygous deletion of the PP2A catalytic subunit α (Ppp2cα) was achieved using Myh6-Cre-mediated recombination. Immunoblot analyses confirmed a marked reduction in PP2A-Cα expression, indicating effective suppression of PP2A catalytic activity in cardiomyocytes (Figure 3A). Under basal conditions, cardiomyocyte-specific PP2A-Cα-deficient mice exhibited normal cardiac structure and function, comparable to those of control mice (Ppp2cα^fl/fl^, Ppp2cα^fl/-^ or Myh6-Cre), after tamoxifen administration (Supplementary Figure 3A and 3B). In subsequent studies, Myh6-Cre mice served as the controls. After acute ISO treatment, PP2A-Cα-deficient mice developed significantly more severe TTS-like phenotypes than control mice. Echocardiographic assessment revealed a greater reduction in left ventricular EF and FS in PP2A-Cα-deficient mice after ISO administration (Figure 3B and supplementary Figure 3C). Consistent with this, histological examination revealed more severe myocardial injury in PP2A-Cα-deficient hearts than in controls (Figure 3C). To further substantiate the functional relevance of PP2A inhibition, we next employed a pharmacological approach using LB-100, a selective PP2A inhibitor. Administration of LB-100 effectively suppressed PP2A activity in the heart, as evidenced by reduced methyl-PP2A-C levels without significant alteration of total PP2A-C expression (Figure 3D). Moreover, mice treated with LB-100 before ISO administration exhibited significantly increased acute mortality compared with ISO alone (Figure 3E). Echocardiographic analysis further revealed that pharmacological inhibition of PP2A alone had minimal effects on cardiac function; however, when combined with ISO challenge, LB-100 markedly aggravated TTS-associated cardiac dysfunction (Figure 3F). In addition, histological analyses corroborated these findings, demonstrating more extensive myocardial injury in hearts from LB-100-treated mice following ISO challenge (Figure 3G).

**Figure 3.**
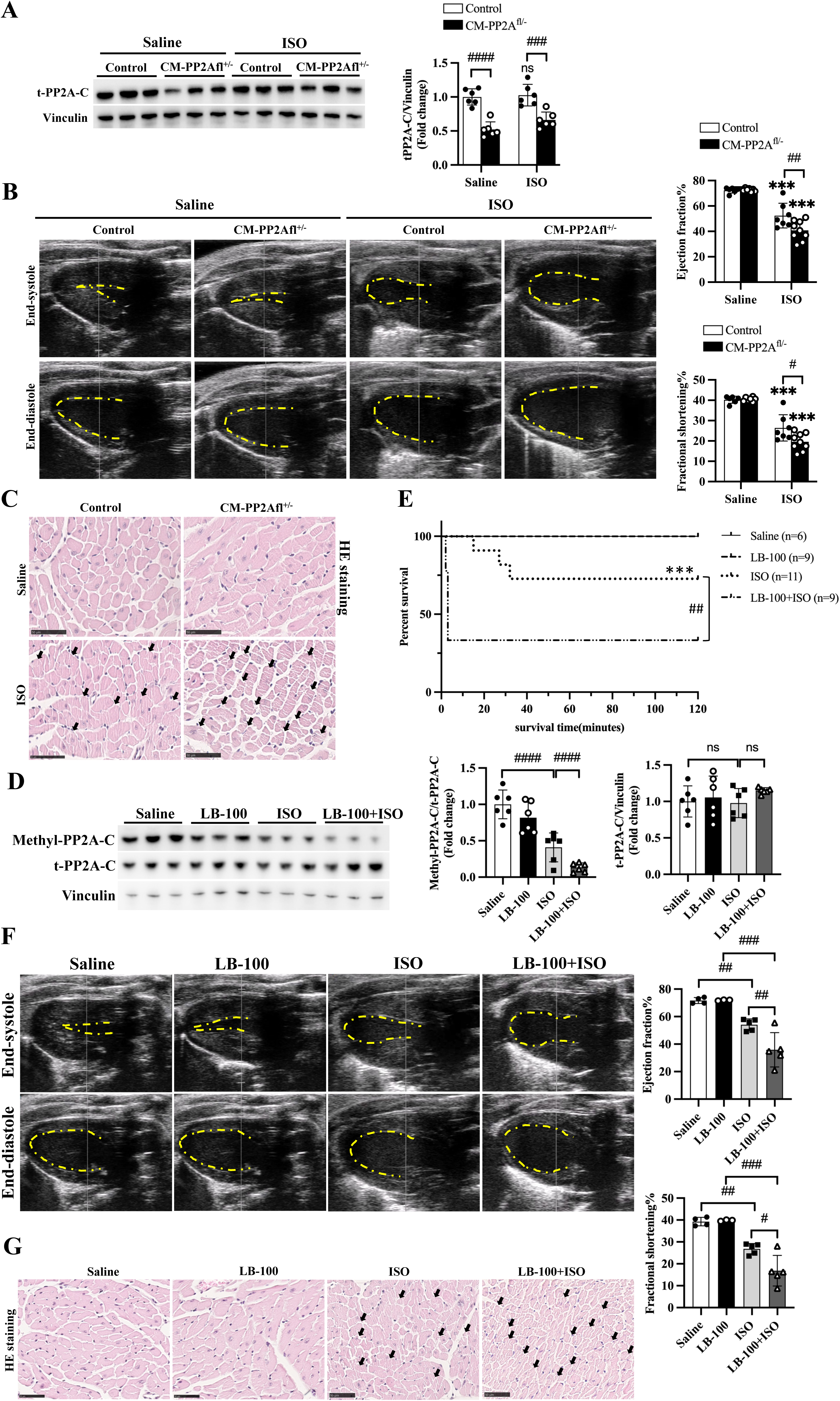
Genetic and pharmacological suppression of PP2A activity exacerbates ISO-induced TTS. A, Immunoblot analysis of total PP2A-C (t-PP2A-C) in cardiac tissue from control (Myh6-Cre) and cardiomyocyte-specific PP2A-Cα–deficient mice (CM-PP2Afl+/-), with quantification normalized to vinculin. **B,** Representative echocardiographic images at end-systole and end-diastole from control and PP2A-Cα–deficient mice under saline or ISO treatment, with quantification of left ventricular EF and FS. **C,** HE staining of cardiac sections from control and PP2A-Cα–deficient mice under saline or ISO treatment. Arrows indicate areas of myocardial injury. **D,** Immunoblot analysis of methyl-PP2A-C and total PP2A-C in cardiac tissue following pretreatment with LB-100 in TTS mcie, with quantification normalized to vinculin. **E,** Kaplan–Meier survival analysis of TTS mice pretreated with LB-100. **F,** Representative echocardiographic images from TTS mice pretreated with LB-100, with quantification of EF and FS. **G,** HE staining of cardiac sections from TTS mice pretreated with LB-100. Arrows indicate areas of myocardial injury. Data are presented as mean±SEM. \**P*<0.05; \*\**P*<0.01; \*\*\**P*<0.001; #*P*<0.05; ##*P*<0.01; ###*P*<0.001; ####*P*<0.0001; ns, not significant.

These genetic and pharmacological data demonstrate that suppression of PP2A activity not only correlates with but also functionally contributes to the severity of acute TTS. Reduced PP2A activity sensitizes the heart to catecholamine-induced stress, thereby exacerbating cardiac dysfunction and myocardial injury during the acute phase of TTS.

### Pharmacological activation of PP2A mitigates acute cardiac dysfunction in TTS

Given that genetic and pharmacological suppression of PP2A activity exacerbated ISO-induced TTS, we next asked whether restoring PP2A activity could protect against acute stress-induced cardiac injury. To this end, mice were treated with DT-061, a selective small-molecule activator of PP2A^32^, after ISO administration (Figure 4A). Functionally, DT-061-mediated PP2A activation significantly improved survival following acute ISO challenge (Figure 4B). Echocardiographic assessment further demonstrated that DT-061 preserved left ventricular systolic function in ISO-treated mice, as evidenced by higher EF and FS compared with vehicle-treated counterparts (Figure 4C). DT-061 treatment alone did not affect cardiac function under basal conditions. Histological examination further showed that DT-061 attenuated ISO-induced acute myocardial injury, with reduced cardiomyocyte damage and preservation of myocardial architecture (Figure 4D). Consistent with these functional and phenotypic improvements, immunohistochemical analysis revealed restoration of methyl-PP2A-C levels in cardiomyocytes from DT-061-treated mice subjected to ISO administration (Figure 4E). Meanwhile, immunoblot analyses confirmed that DT-061 treatment increased methyl-PP2A-C levels in the ISO-treated heart without significantly altering total PP2A-C (Figure 4A), consistent with selective enhancement of PP2A enzymatic activity.

**Figure 4.**
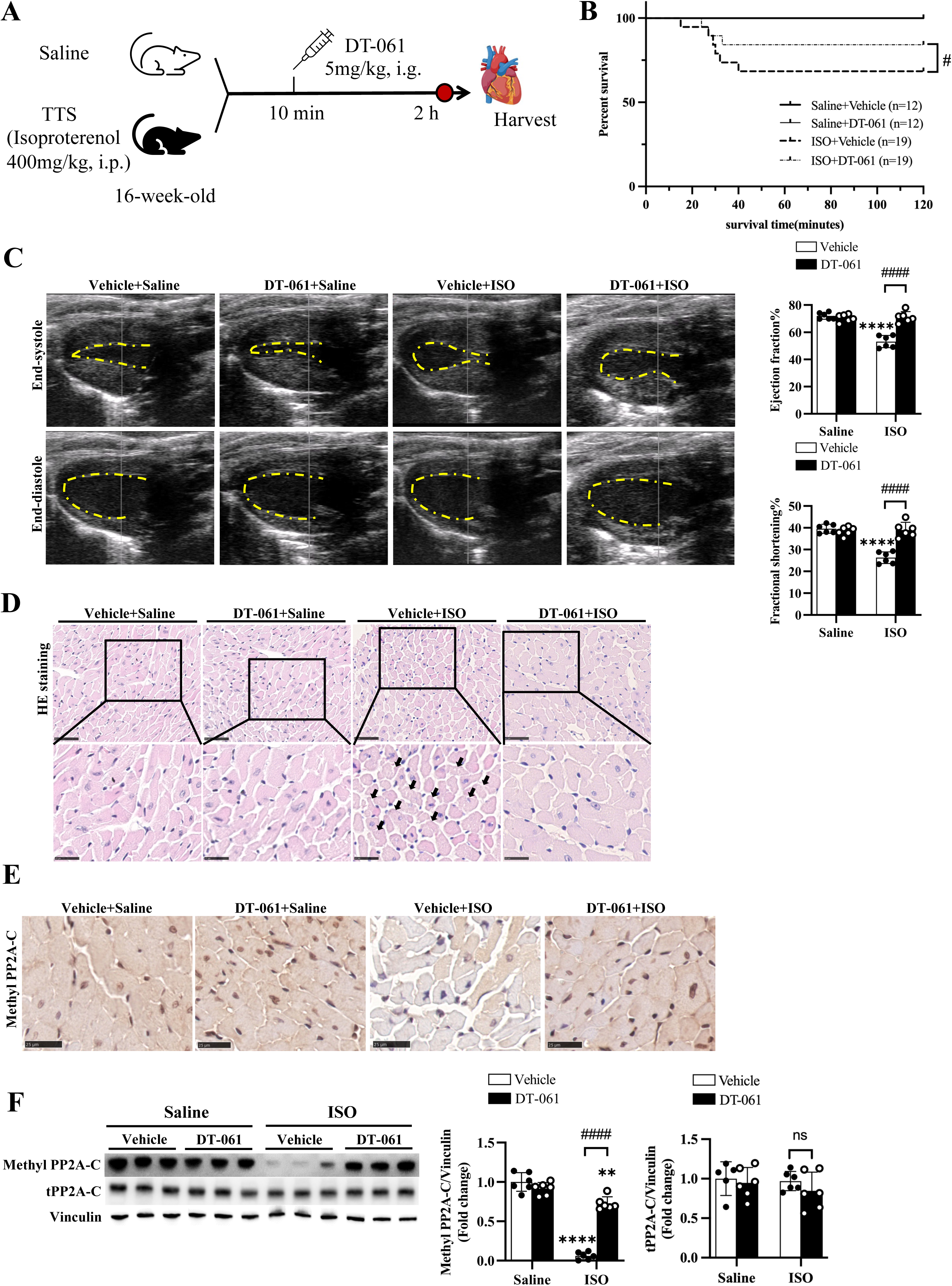
Pharmacological activation of PP2A attenuates acute cardiac dysfunction and myocardial injury in TTS. A, Schematic diagram showing the mouse model of TTS followed by treatment with the PP2A activator DT-061. **B,** Kaplan–Meier survival analysis of TTS mice followed by treatment with DT-061. **C,** Representative echocardiographic images at end-systole and end-diastole from TTS mice followed by treatment with DT-061, with quantification of left ventricular EF and FS. **D,** HE staining of cardiac sections from the indicated groups. Arrows indicate areas of myocardial injury. **E,** Immunohistochemical staining of methyl-PP2A-C in cardiac tissue from the indicated groups. **F,** Immunoblot analysis of methyl-PP2A-C and total PP2A-C (t-PP2A-C) in cardiac tissue from the indicated groups, with quantification normalized to vinculin. Data are presented as mean±SEM. \**P*<0.05; \*\**P*<0.01; \*\*\**P*<0.001; \*\*\*\**P*<0.0001; #*P*<0.05; ##*P*<0.01; ###*P*<0.001; ####*P*<0.0001; ns, not significant.

To assess whether the protective effects of PP2A activation are conserved across distinct experimental models of TTS, we next used an epinephrine (EPI)-induced TTS model.^33^ Consistent with observations in the ISO model, EPI administration markedly reduced methyl-PP2A-C levels in the heart, indicating suppression of PP2A activity, whereas total PP2A-C expression remained unchanged (Supplementary Figure 4A). Importantly, DT-061 treatment restored methyl-PP2A-C levels in EPI-treated mice without altering total PP2A-C abundance, consistent with selective rescue of PP2A activity. Functionally, EPI-treated mice developed significant left ventricular systolic dysfunction, which was substantially ameliorated by DT-061 treatment, as evidenced by improved echocardiographic parameters (Supplementary Figure 4B). In parallel, histological analyses revealed that DT-061 attenuated myocardial injury in the EPI-induced TTS model (Supplementary Figure 4C).

Taken together, these data demonstrate that pharmacological activation of PP2A significantly mitigates acute cardiac dysfunction and myocardial injury in multiple experimental models of Takotsubo syndrome, underscoring PP2A activity as a key modulator of disease severity during the acute phase of TTS.

### PP2A activity modulates ferritinophagy-mediated ferroptosis and mitochondrial injury in TTS

To elucidate downstream cellular processes regulated by PP2A during TTS, we performed bulk RNA sequencing of cardiac tissue from ISO-induced TTS mice treated with vehicle or DT-061. Unsupervised hierarchical clustering of differentially expressed genes revealed a distinct transcriptional signature in ISO-treated hearts compared with saline controls, which was partially normalized by DT-061 treatment (Figure 5A). Volcano plot analysis further highlighted the magnitude and directionality of these changes. Relative to saline controls, ISO challenge induced widespread transcriptional remodeling, with a predominance of upregulated genes consistent with activation of a coordinated stress-responsive program (Figure 5B, left). In contrast, comparison of DT-061-treated versus vehicle-treated ISO hearts revealed extensive bidirectional gene regulation (Figure 5B, right), with a substantial subset of ISO-upregulated genes suppressed following PP2A activation and many ISO-downregulated genes partially restored.

**Figure 5.**
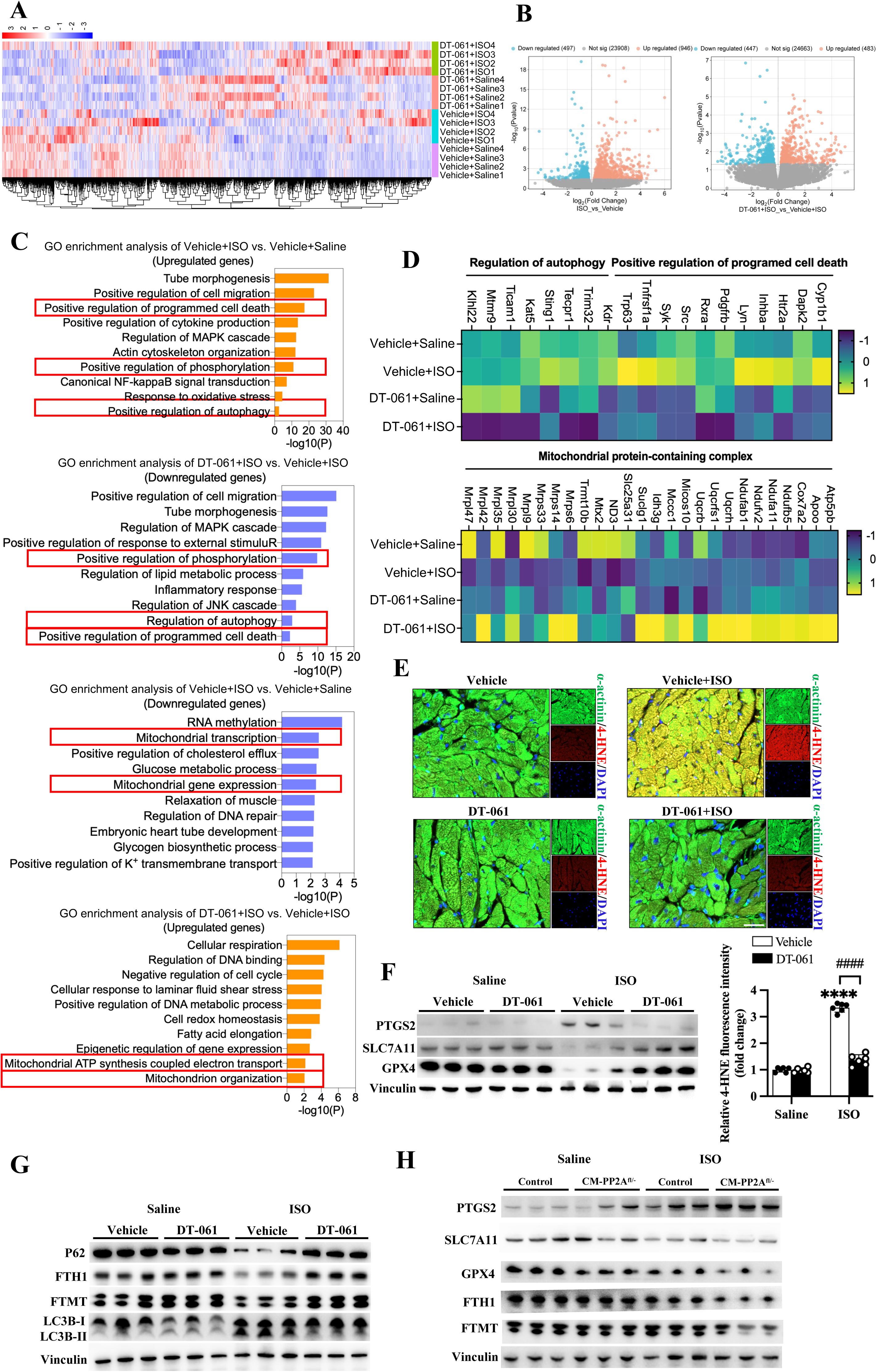
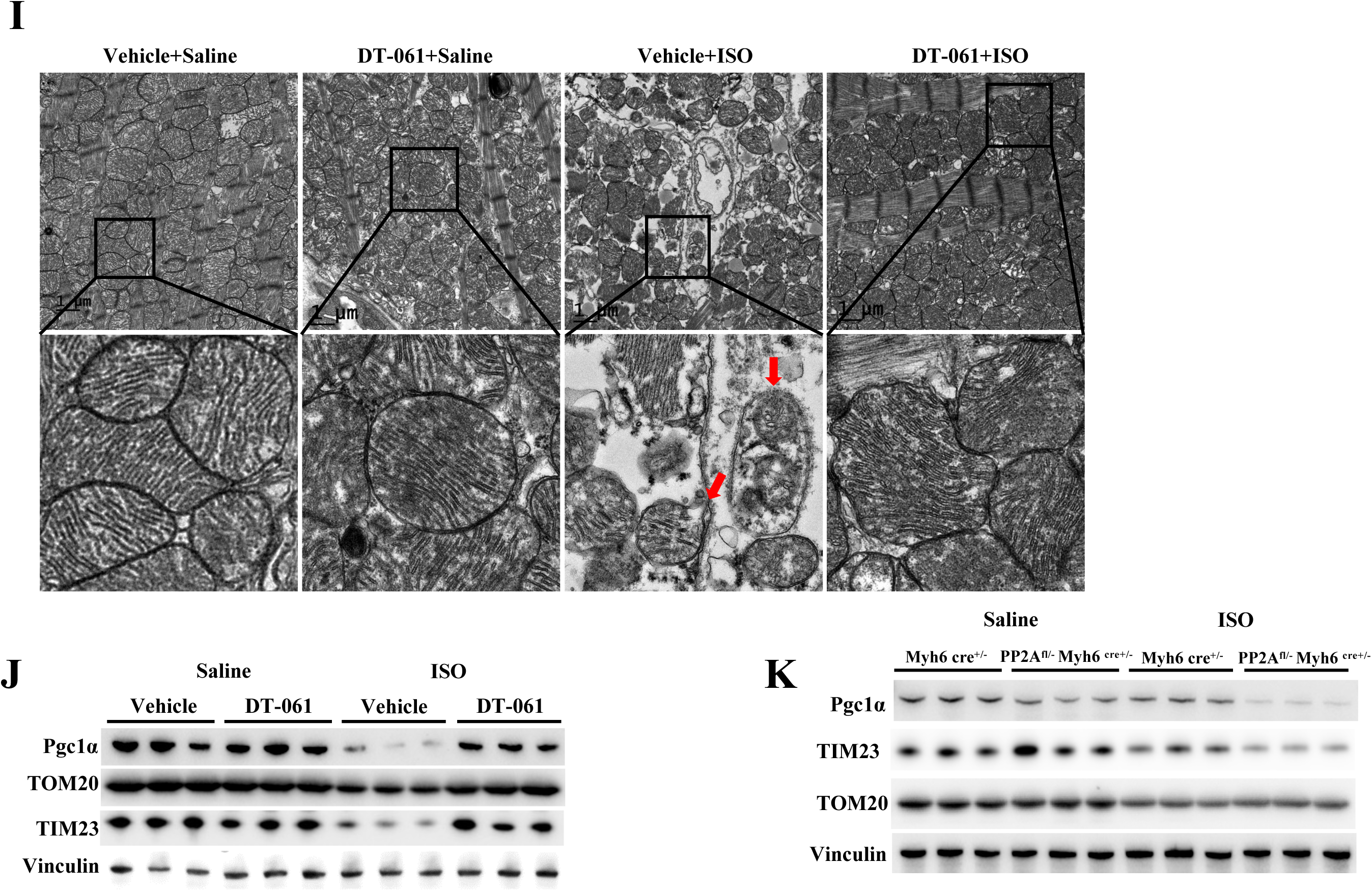
PP2A activity modulates ferritinophagy-associated ferroptosis and mitochondrial injury in TTS. A, Unsupervised hierarchical clustering heatmap of differentially expressed genes in cardiac tissue from DT-061-treated TTS mice. **B,** Volcano plots showing differential gene expression between saline and ISO groups (left), and between vehicle and DT-061 groups after ISO treatment (right). **C,** GO enrichment analysis of ISO-induced genes followed by DT-061 treatment. **D,** Heatmaps of representative genes of of ISO-induced TTS mice followed by DT-061 treatment. **E,** Representative immunofluorescence images of 4-hydroxynonenal (4-HNE) staining in cardiac tissue. **F,** Immunoblot analysis of ferroptosis-associated markers in cardiac tissue from the indicated groups. **G,** Immunoblot analysis of ferritinophagy-related proteins in cardiac tissue from the indicated groups. **H,** Immunoblot analysis of ferritinophagy-related proteins in cardiomyocyte-specific PP2A-Cα–deficient mice. **I,** Transmission electron microscopy images showing mitochondrial ultrastructure and autophagic structures in the indicated groups. Arrows indicate damaged mitochondria or autophagic vacuoles. **J,** Immunoblot analysis of mitochondrial proteins in cardiac tissue from the indicated groups. **K,** Immunoblot analysis of mitochondrial proteins in cardiomyocyte-specific PP2A-Cα–deficient mice. Data are presented as mean±SEM. Data are presented as mean±SEM. \**P*<0.05; \*\**P*<0.01; \*\*\**P*<0.001; \*\*\*\**P*<0.0001; #*P*<0.05; ##*P*<0.01; ###*P*<0.001; ####*P*<0.0001; ns, not significant.

Functional enrichment analysis of ISO-upregulated genes demonstrated significant overrepresentation of pathways related to regulated cell death, autophagy, and mitochondrial processes. Notably, enrichment of these stress-associated pathways was attenuated in DT-061-treated hearts, suggesting that PP2A activation broadly dampens ISO-induced transcriptional stress responses (Figure 5C). Consistent with these pathway-level findings, heatmap analysis of representative genes showed coordinated induction following ISO exposure and partial reversal with DT-061 treatment (Figure 5D). Together, these data indicate that acute TTS is characterized by activation of cell death-, autophagy-, and mitochondrial-associated gene programs, which are globally modulated by PP2A activity.

Guided by these transcriptomic findings, we next assessed major forms of programmed cell death at the protein level. Immunoblot analyses showed that canonical markers of apoptosis and pyroptosis were not significantly altered following ISO challenge (Supplementary Figure 5A). In contrast, immunofluorescence revealed marked myocardial lipid peroxidation, as evidenced by robust accumulation of 4-hydroxynonenal (4-HNE) in cardiomyocytes, which was substantially attenuated by DT-061 treatment (Figure 5E). Consistently, immunoblotting demonstrated a ferroptosis-associated signature in ISO-treated hearts, characterized by increased PTGS2 expression and reduced levels of the antioxidant regulators GPX4 and SLC7A11, all of which were largely normalized by DT-061 activation (Figure 5F and Supplementary Figure 6A). Given that transcriptomic analysis also suggested enhanced autophagic activity, we next examined ferritinophagy, a selective autophagy pathway involved in intracellular iron handling. ISO-induced TTS hearts exhibited reduced levels of cytosolic ferritin heavy chain (FTH1) and mitochondrial ferritin (FTMT), accompanied by decreased p62 and an increased LC3B-II/LC3B-I ratio, consistent with enhanced ferritinophagy. Importantly, activation of PP2A by DT-061 largely reversed these changes, indicating that PP2A activity restrains excessive ferritinophagy during acute stress (Figure 5G and Supplementary Figure 6B). Conversely, cardiomyocyte-specific PP2A-Cα knockdown further exacerbated ISO-induced ferritinophagy-related alterations, supporting a regulatory role for PP2A in iron-associated autophagic processes (Figure 5H and Supplementary Figure 7A).

In parallel with altered iron handling, ISO-induced TTS was associated with pronounced mitochondrial abnormalities. Transmission electron microscopy revealed increased autophagic structures and marked mitochondrial ultrastructural disruption, including swelling and loss of outer membrane integrity, in ISO-treated hearts. These abnormalities were substantially alleviated by DT-061 administration (Figure 5I). Consistently, ISO challenge induced dysregulation of key mitochondrial regulators and structural proteins, including PGC1α, TIM23, and TOM20, whereas PP2A activation preserved mitochondrial protein homeostasis (Figure 5J and Supplementary Figure 6C). In contrast, cardiomyocyte-specific PP2A-Cα deficiency further exacerbated these mitochondrial alterations (Figure 5K and Supplementary Figure 7B).

To assess the generalizability of these findings, we evaluated an independent TTS model induced by EPI. Consistent with the ISO model, EPI challenge induced ferroptosis-associated changes, enhanced ferritinophagy, and mitochondrial injury, several of which were attenuated by DT-061 treatment (Supplementary Figure 8A and 8B).

Collectively, these results demonstrate that acute TTS is characterized by concurrent activation of ferritinophagy-associated ferroptosis and mitochondrial dysfunction, which are dynamically modulated by PP2A activity. Dysregulation of these interconnected stress-response pathways likely contributes to cardiomyocyte vulnerability during acute catecholamine stress.

### PP2A activity regulates ferroptosis and mitochondrial dysfunction in cardiomyocytes in vitro

To establish an in vitro model of PP2A regulation under catecholamine stress, we first examined methylated PP2A-C expression in cardiomyocytes. Immunofluorescence confirmed robust basal expression of methyl-PP2A-C in both H9C2 cells and primary human cardiomyocytes (Supplementary Figures 9A and 11A). To define appropriate experimental conditions, H9C2 cells were exposed to increasing ISO concentrations (1, 5, and 10 mM). Immunoblot analysis demonstrated a dose-dependent reduction in methyl-PP2A-C levels, with a consistent, pronounced decrease at 10 mM ISO (Supplementary Figure 9B). This concentration was therefore used for subsequent experiments. Under these conditions, ISO exposure significantly increased cardiomyocyte death, as assessed by cell death staining. This effect was markedly attenuated by treatment with compound DT-061, a PP2A-C activator, whereas pharmacologic inhibition of PP2A-C with LB-100 further exacerbated ISO-induced cell death (Figure 6A). Given the established link between ferroptosis and lipid peroxidation, we next evaluated lipid reactive oxygen species (ROS) accumulation. Fluorescent lipid ROS staining revealed a substantial increase in lipid peroxidation following ISO treatment, which was significantly reduced by DT-061 and further enhanced by LB-100 (Figure 6B).

**Figure 6.**
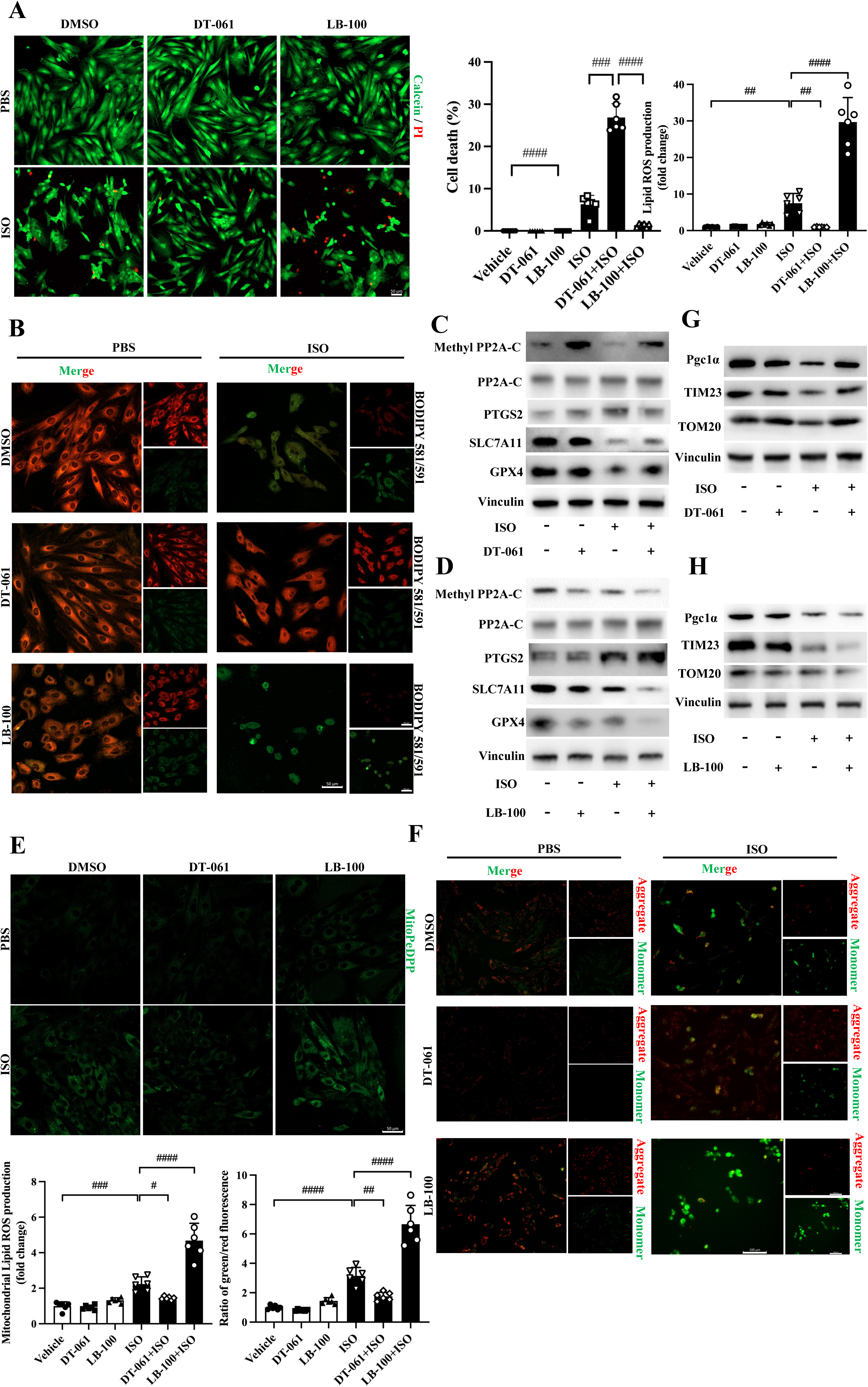
PP2A activity regulates ferroptosis and mitochondrial dysfunction in cardiomyocytes in vitro. A, Representative images of calcein-AM/propidium iodide (PI) staining in H9C2 cells treated with DT-061 or LB-100 under basal (PBS) or ISO conditions, with quantification of cell death. **B,** Representative fluorescence images of lipid reactive oxygen species (ROS) using BODIPY 581/591 C11 staining in H9C2 cells under the indicated conditions, with quantification of lipid ROS production. **C,** Immunoblot analysis of methyl-PP2A-C, t-PP2A-C, and ferroptosis-associated proteins (PTGS2, SLC7A11, GPX4) in H9C2 cells treated with ISO in the presence or absence of DT-061. **D,** Immunoblot analysis of methyl-PP2A-C, t-PP2A-C, and ferroptosis-associated proteins in H9C2 cells treated with ISO in the presence or absence of LB-100. **E,** Mitochondrial lipid ROS assessed by MitoPeDPP staining in H9C2 cells under the indicated conditions, with quantification. **F,** Assessment of mitochondrial membrane potential using JC-1 staining in H9C2 cells under the indicated conditions, shown as representative images and quantification of red-to-green fluorescence ratio. **G,** Immunoblot analysis of mitochondrial regulatory proteins (PGC1α, TIM23, TOM20) in H9C2 cells treated with ISO in the presence or absence of DT-061. **H,** Immunoblot analysis of mitochondrial proteins in H9C2 cells treated with ISO in the presence or absence of LB-100. Data are presented as mean±SEM. #*P*<0.05; ##*P*<0.01; ###*P*<0.001; ####*P*<0.0001; ns, not significant.

At the molecular level, immunoblot analyses confirmed that DT-061 increased methyl-PP2A-C levels and mitigated the ISO-induced reduction in methyl-PP2A-C, consistent with enhanced PP2A enzymatic activity (Figure 6C; Supplementary Figure 10A). Correspondingly, DT-061 attenuated ISO-induced ferroptosis-associated molecular changes, including reduced PTGS2 expression and restoration of the antioxidant proteins SLC7A11 and GPX4 (Figure 6C; Supplementary Figure 10A). Notably, these protective effects were recapitulated in primary human cardiomyocytes, supporting a conserved anti-ferroptotic role of PP2A across models (Supplementary Figures 11B and 11D). In contrast, pharmacological inhibition of PP2A with LB-100 decreased methyl-PP2A-C levels at baseline and further suppressed methyl-PP2A-C following ISO treatment in both H9C2 cells and primary human cardiomyocytes (Figure 6D; Supplementary Figures 10B and 11C). This reduction in PP2A activity was accompanied by exacerbation of ferroptosis-associated protein changes across both models (Figure 6D; Supplementary Figure 10B and 11E). Together, these findings indicate that PP2A activity inversely regulates ferroptosis in cardiomyocytes, with activation conferring protection and inhibition exacerbating injury under catecholamine stress.

We next assessed mitochondrial lipid peroxidation using MitoPeDPP staining showed a similar pattern, with PP2A-C activation suppressing and PP2A inhibition exacerbating ISO-induced mitochondrial lipid ROS accumulation (Figure 6E). Moreover, mitochondrial function was assessed using JC-1 staining. The results showed ISO exposure induced a marked loss of mitochondrial membrane potential, reflected by a decreased red-to-green fluorescence ratio in H9C2 cells. PP2A activation partially preserved mitochondrial membrane potential, whereas PP2A inhibition further aggravated mitochondrial depolarization (Figure 6F). Consistent with these functional data, ISO-induced dysregulation of key mitochondrial proteins, including PGC1α, TIM23, and TOM20, was attenuated by DT-061and exacerbated by LB-100 in both H9C2 cells and primary human cardiomyocytes (Figure 6G and 6H; Supplementary Figures 10C and 10D; Supplementary Figures 11F and 11G).

Collectively, these in vitro findings demonstrate that PP2A activity modulates ferroptosis-associated oxidative stress and mitochondrial dysfunction in cardiomyocytes, providing mechanistic support for the in vivo observations in experimental models of TTS.

### PP2A modulates ferritinophagy-dependent ferroptosis and mitochondrial dysfunction in cardiomyocytes

To further define the contribution of iron handling and ferritinophagy to PP2A activity loss-regulated cardiomyocyte injury, we examined intracellular iron dynamics in vitro. Cytosolic ferrous iron (Fe²□) was assessed using FerroOrange staining. ISO exposure induced a marked increase in cytosolic Fe²□ levels, which was further exacerbated by pharmacological inhibition of PP2A with LB-100, whereas activation of PP2A with DT-061 significantly attenuated iron accumulation in H9C2 cardiomyocytes (Figure 7A).

**Figure 7.**
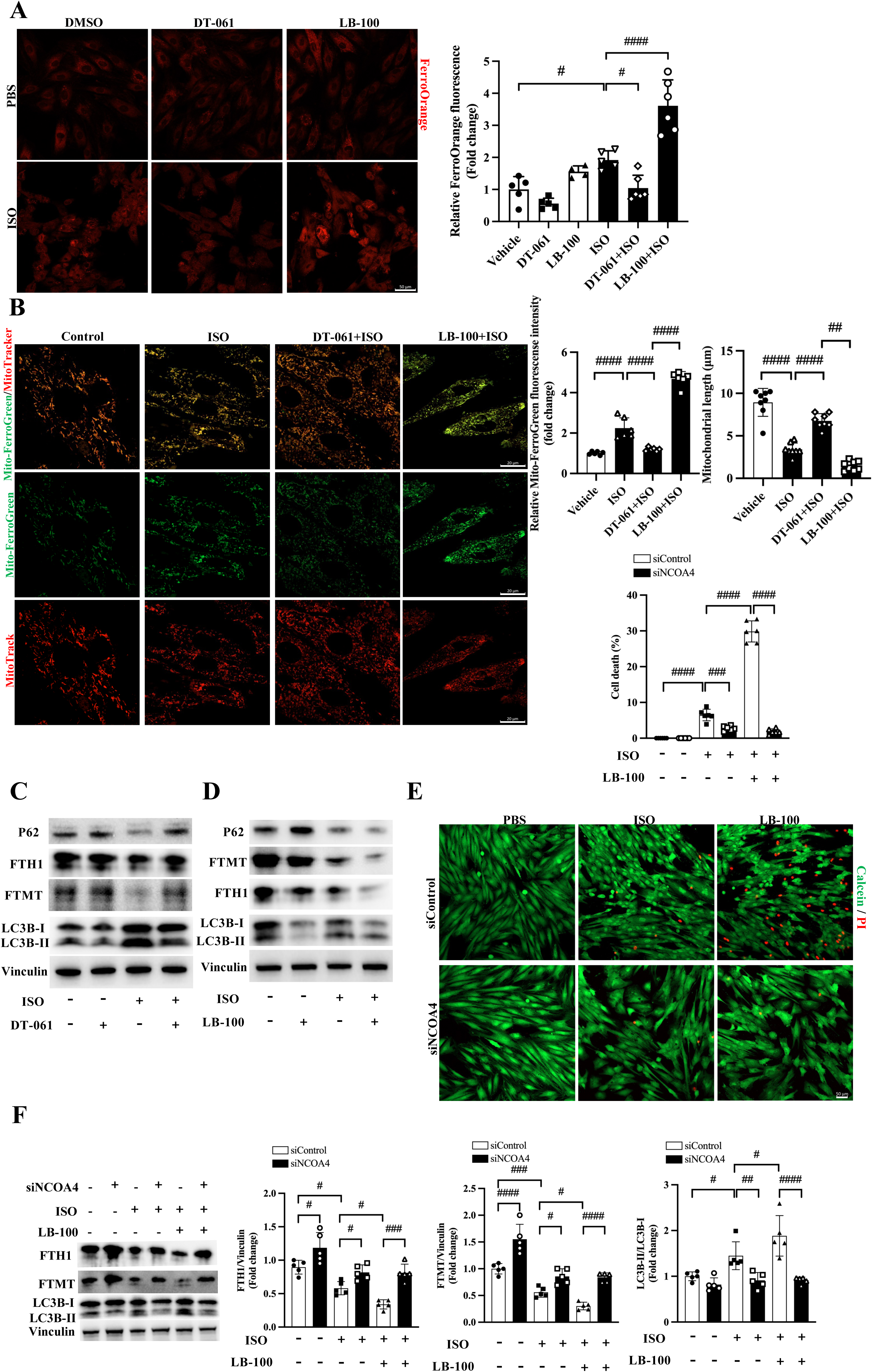
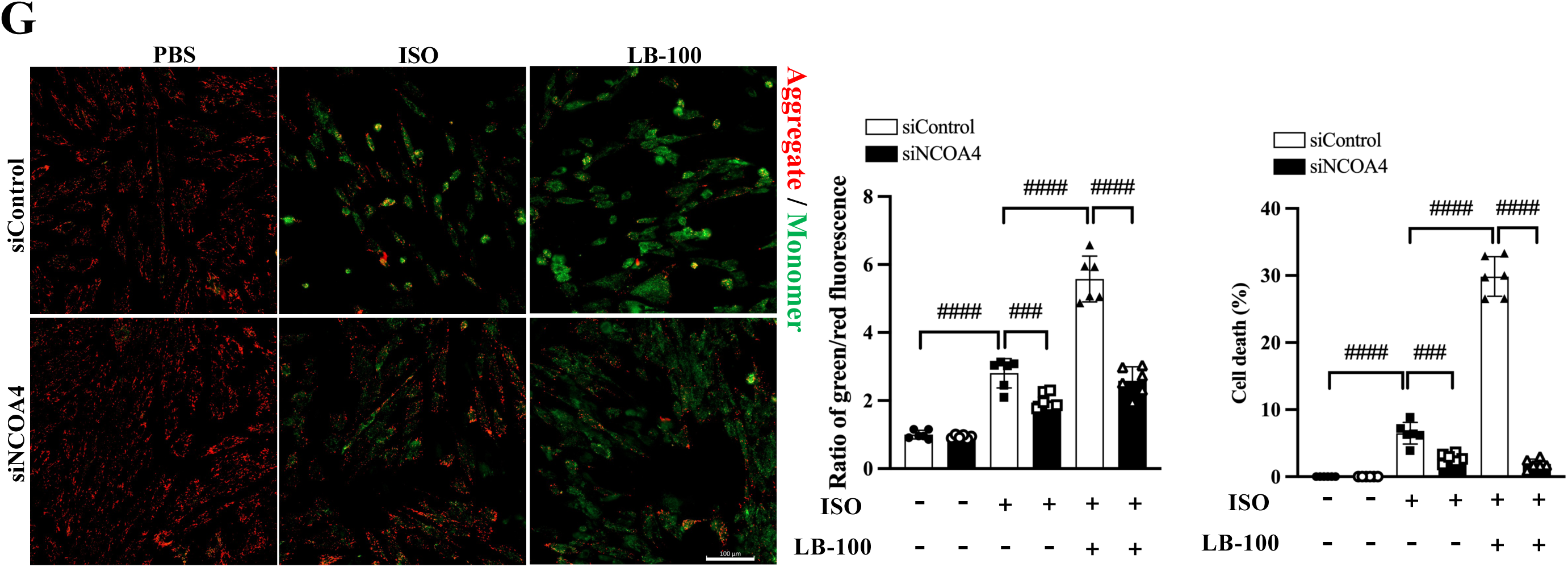
PP2A regulates ferritinophagy-dependent iron homeostasis and mitochondrial integrity in cardiomyocytes. A, Representative images and quantification of cytosolic ferrous iron (Fe²⁺) measured by FerroOrange staining in H9C2 cells treated with DT-061 or LB-100 under ISO administration. **B,** Representative confocal images of mitochondrial iron (Mito-FerroGreen) and mitochondrial morphology (MitoTracker) in H9C2 cells under the indicated conditions, with quantification of mitochondrial Fe²⁺ levels and mitochondrial length. **C,** Immunoblot analysis of ferritinophagy-related proteins in H9C2 cells treated with ISO in the presence or absence of DT-061. **D,** Immunoblot analysis of ferritinophagy-related proteins in H9C2 cells treated with ISO in the presence or absence of LB-100. **E,** Representative images and quantification of cardiomyocyte death following NCOA4 knockdown under ISO and LB-100 treatment. **F,** Immunoblot analysis of FTH1 and FTMT following NCOA4 silencing in H9C2 cells. **G,** Assessment of mitochondrial membrane potential by JC-1 staining following NCOA4 knockdown, with quantification of red-to-green fluorescence ratio. Data are presented as mean±SEM. *P<0.05; **P<0.01; ***P<0.001; ****P<0.0001; #P<0.05; ##P<0.01; ###P<0.001; ####P<0.0001; ns, not significant.

To determine the subcellular distribution of iron, we performed confocal co-staining with MitoTracker. FerroOrange fluorescence exhibited minimal overlap with mitochondrial labeling, indicating that the increased signal primarily reflects expansion of the cytosolic labile Fe²□ pool in both H9C2 and primary human cardiomyocytes (Supplementary Figures 12A and 13A). Mitochondrial iron was therefore assessed independently using Mito-FerroGreen. ISO treatment induced a pronounced increase in mitochondrial Fe²□ accumulation, which was further enhanced by PP2A inhibition and significantly reduced by PP2A activation across both cardiomyocyte models (Figure 7B; Supplementary Figure 13B). In parallel, mitochondrial imaging revealed marked structural abnormalities following ISO exposure, including fragmentation and disruption of mitochondrial networks; these changes were aggravated by LB-100 and partially preserved by DT-061 treatment (Figure 7B and Supplementary Figure 13B).

At the molecular level, immunoblot analyses showed that ISO-induced stress reduced levels of cytosolic ferritin heavy chain (FTH1) and mitochondrial ferritin (FTMT), consistent with increased ferritin turnover. These effects were exacerbated by PP2A inhibition and attenuated by PP2A activation in both H9C2 cells and primary human cardiomyocytes (Figure 7C and 7D; Supplementary Figures 12B and 12C; Supplementary Figures 13C and 13D), indicating that PP2A activity restrains excessive ferritin degradation during catecholamine stress. To determine whether this process is autophagy-dependent, cells were treated with the lysosomal inhibitor bafilomycin A1. As expected, bafilomycin A1 led to accumulation of LC3-II and p62, confirming effective blockade of autophagic flux. Under these conditions, degradation of FTH1 and FTMT induced by ISO and LB-100 was partially prevented, supporting an autophagy-dependent mechanism for ferritin turnover in PP2A-regulated iron homeostasis (Supplementary Figure 12D).

To further assess the functional importance of mitochondrial ferritin, FTMT was silenced in H9C2 cardiomyocytes. FTMT knockdown induced pronounced mitochondrial fragmentation and structural disorganization even under basal conditions. Notably, FTMT silencing abrogated the protective effects of PP2A activation on mitochondrial morphology following ISO exposure, resulting in persistent mitochondrial disruption despite DT-061 treatment (Supplementary Figure 12E). These findings identify mitochondrial ferritin as a critical determinant of mitochondrial integrity and suggest that its depletion represents a key downstream event linking dysregulated iron handling to mitochondrial injury.

Because nuclear receptor coactivator 4 (NCOA4) is a key cargo receptor that mediates ferritinophagy, we next disrupted this pathway using siRNA. Importantly, NCOA4 knockdown significantly reduced cardiomyocyte death induced by combined ISO exposure and PP2A inhibition (Figure 7E). In addition, NCOA4 silencing attenuated LB-100–enhanced degradation of FTH1 and FTMT, confirming effective inhibition of ferritinophagy (Figure 7F). Consistent with this reversal, JC-1 staining demonstrated partial preservation of mitochondrial membrane potential after NCOA4 silencing (Figure 7G).

Collectively, these data demonstrate that PP2A activity regulates ferritinophagy-dependent ferroptotic stress in cardiomyocytes. Dysregulated ferritinophagy promotes the accumulation of cytosolic and mitochondrial iron, leading to mitochondrial dysfunction and increased cell death under catecholamine stress. Restoring PP2A activity mitigates these pathological processes.

### PP2A restrains ferritinophagy and mitochondrial injury by suppressing JNK-MAPK signaling in TTS

As shown above, comparative transcriptomic profiling of cardiac tissue identified MAPK signaling as a major stress-responsive pathway in ISO-induced TTS and showed marked suppression following PP2A activation with compound DT-061 (Figure 5C). We next investigated whether MAPK signaling contributes to PP2A-mediated cardioprotection during acute stress. Immunoblot analysis demonstrated that JNK phosphorylation was robustly increased following ISO challenge and significantly suppressed by DT-061 treatment, whereas ERK and p38 phosphorylation were not appreciably altered (Figure 8A; Supplementary Figure 14A). These findings identify JNK as the predominant MAPK pathway activated during acute TTS and regulated by PP2A activity.

**Figure 8.**
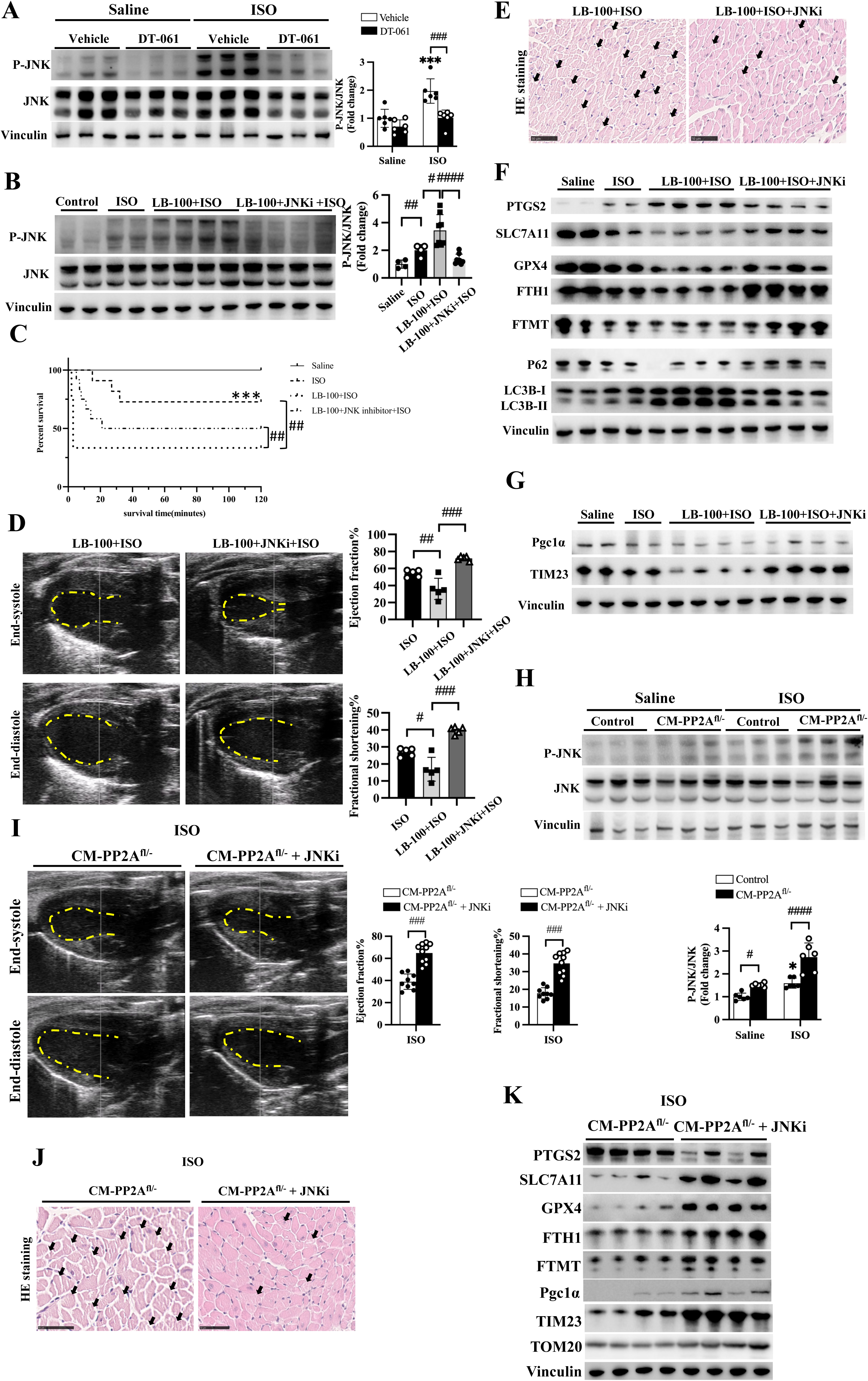
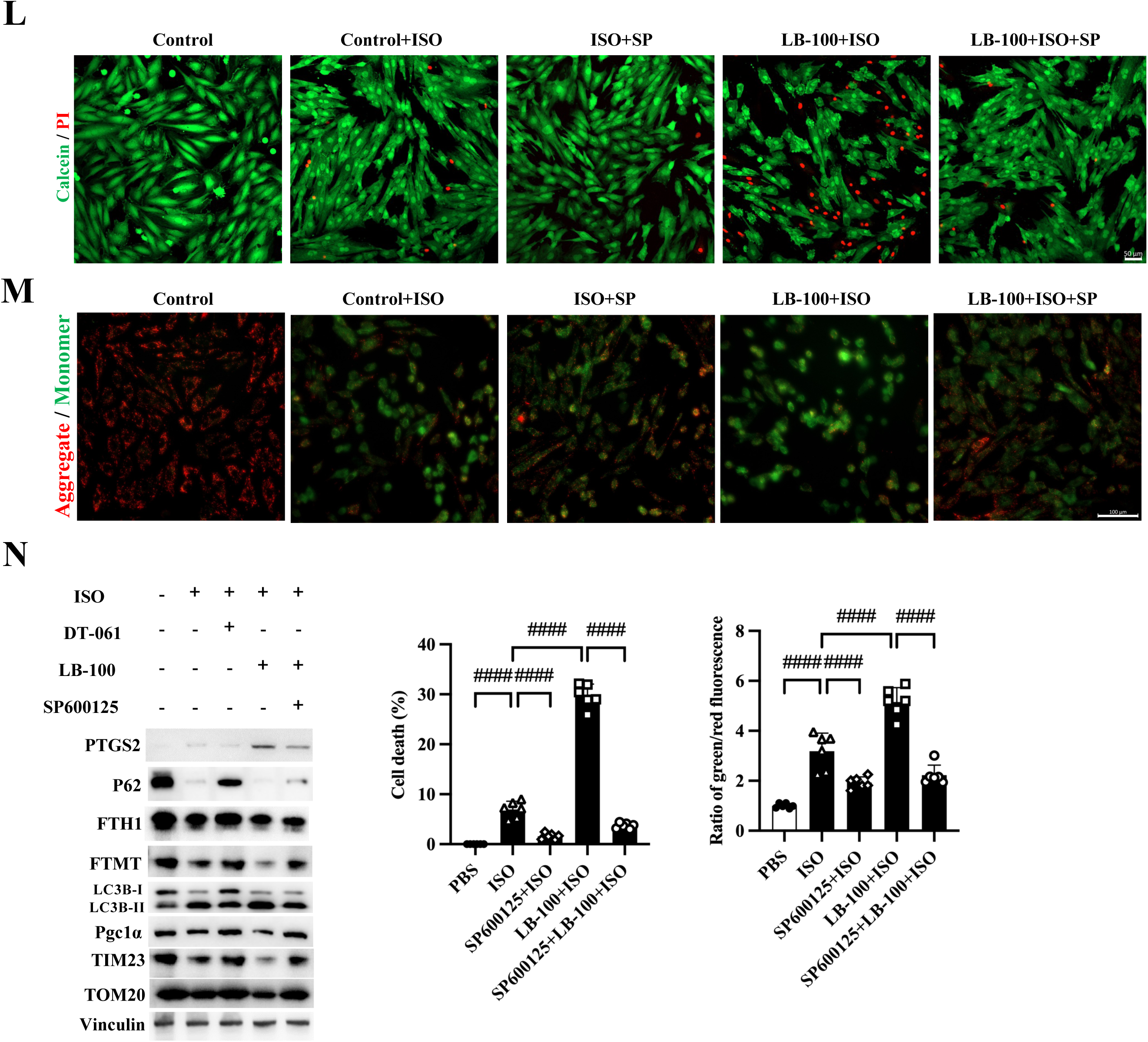
PP2A restrains ferritinophagy and mitochondrial injury via suppression of JNK signaling in TTS. A, Immunoblot analysis and quantification of JNK pathway in mouse hearts following ISO treatment with or without DT-061. **B,** Immunoblot analysis and quantification of JNK pathway in mouse hearts following ISO treatment with or without LB-100 or JNK inhibitor. **C,** Kaplan– Meier survival curves of mice treated with ISO and LB-100 in the presence or absence of the JNK inhibitor SP600125. **D,** Representative echocardiographic and quantification of left ventricular EF and FS in mice subjected to ISO and LB-100 with or without SP600125. **E,** Representative HE staining of myocardial sections showing cardiac injury under the indicated conditions. **F,** Immunoblot analysis of ferritinophagy- and ferroptosis-related proteins in mouse hearts following JNK inhibition. **G,** Immunoblot analysis of mitochondrial regulatory proteins following JNK inhibition. **H,** Immunoblot analysis of JNK phosphorylation in cardiomyocyte-specific PP2A-Cα–deficient mice treated with ISO. **I,** Representative echocardiographic and quantification of left ventricular EF and FS in PP2A-Cα– deficient TTS mice with JNK inhibition. **J,** Representative HE staining showing myocardial injury in PP2A-Cα–deficient TTS mice with or without JNK inhibition. **K,** Immunoblot analysis of ferritinophagy- and mitochondrial-associated proteins in PP2A-Cα–deficient TTS mice following JNK inhibition. **L,** Representative images and quantification of cardiomyocyte death in H9C2 cells treated with ISO and LB-100 in the presence or absence of SP600125. **M,** Assessment of mitochondrial membrane potential in H9C2 cells under the indicated conditions. **N,** Immunoblot analysis of ferritinophagy- and mitochondrial-related proteins in H9C2 cells following JNK inhibition. Data are presented as mean±SEM. *P<0.05; **P<0.01; ***P<0.001; ****P<0.0001; #P<0.05; ##P<0.01; ###P<0.001; ####P<0.0001; ns, not significant.

To determine the functional contribution of JNK signaling, we pharmacologically inhibited JNK with SP600125 in mice subjected to combined PP2A inhibition and ISO challenge. JNK inhibition effectively reduced elevated p-JNK levels and significantly improved mortality in those mice (Figure 8B and 8C). Moreover, JNK inhibition significantly ameliorated left ventricular systolic dysfunction, as assessed by echocardiography (Figure 8D). Histological analysis further demonstrated attenuation of ISO-induced myocardial injury that was exacerbated by PP2A inhibition (Figure 8E).

At the molecular level, JNK inhibition substantially reversed LB-100–exacerbated alterations in ferritinophagy-associated markers, including restoring ferritin levels and attenuating ferroptosis-related proteins (Figure 8F and Supplementary Figure 16A). In parallel, JNK inhibition significantly normalized dysregulation of mitochondrial regulatory and structural proteins induced by combined PP2A inhibition and ISO challenge (Figure 8G and Supplementary Figure 16B), indicating that JNK activation mediates mitochondrial injury associated with impaired PP2A activity during TTS.

To validate these findings in a genetic model, cardiomyocyte-specific PP2A-Cα-deficient mice were treated with SP600125 prior to ISO challenge. Consistent with the pharmacological model, JNK inhibition suppressed elevated p-JNK levels in PP2A-Cα-deficient hearts following ISO administration (Figure 8H). Functionally, JNK inhibition markedly improved cardiac performance, as evidenced by recovery of left ventricular systolic function (Figure 8I) and reduced myocardial injury on histological assessment (Figure 8J). Immunoblot analysis further confirmed that JNK inhibition attenuated PP2A-Cα-deficiency-induced ferritinophagy-associated changes and mitochondrial dysregulation (Figure 8K and Supplementary Figure 16C).

We next examined whether JNK signaling similarly mediates PP2A-dependent protection in vitro. In H9C2 cardiomyocytes, ISO-induced JNK phosphorylation was further enhanced by PP2A inhibition and attenuated by PP2A activation (Supplementary Figure 15A and 15B). JNK inhibition significantly reduced cardiomyocyte death and lipid peroxidation during combined ISO exposure and PP2A inhibition (Figure 8L; Supplementary Figure 15C). Consistent with this, JC-1 and MitoTracker staining demonstrated partial preservation of mitochondrial membrane potential and morphology following JNK inhibition (Figure 8M; Supplementary Figure 15D). In addition, immunoblot analysis showed that JNK inhibition mitigated PP2A inhibition-induced alterations in ferritinophagy- and mitochondrial-associated proteins during catecholamine stress in vitro(Figure 8N and Supplementary Figure 16D).

Collectively, these findings demonstrate that reduced PP2A activity triggers JNK-MAPK signaling during acute TTS. JNK serves as a key downstream mediator linking impaired PP2A activity to ferritinophagy, ferroptosis, and mitochondrial dysfunction, whereas pharmacological inhibition of JNK mitigates cardiomyocyte injury in both in vivo and in vitro models.

## DISCUSSION

This study identifies a previously unrecognized role for loss of PP2A activity in the pathogenesis of acute TTS. By integrating multi-omics analyses, human patient samples, and experimental models, we show that PP2A activity is rapidly and regionally suppressed during catecholamine stress. Using complementary in vivo loss- and gain-of-function approaches, we demonstrate that PP2A inactivation exacerbates stress-induced cardiac dysfunction and myocardial injury, whereas restoring PP2A activity confers robust cardioprotection. Mechanistically, PP2A inactivation drives JNK–MAPK signaling, leading to dysregulated iron handling via ferritinophagy, mitochondrial dysfunction, and subsequent cardiomyocyte injury. Collectively, these findings position PP2A as a central signaling node linking catecholamine stress to reversible myocardial injury and identify the PP2A–JNK axis as a potential therapeutic target in acute TTS.

Previous studies have implicated a wide range of signaling pathways in the pathogenesis of TTS, including stress-activated kinase cascades, oxidative stress responses, regulated cell death programs, and mitochondrial dysfunction, underscoring the complexity of the molecular processes underlying catecholamine-induced myocardial injury.^30,31,33^ However, these pathways have largely been identified in individual experimental settings or specific disease contexts, and it has remained unclear whether they represent model-specific phenomena or reflect conserved molecular signatures shared across TTS. Addressing this gap, our integrative analyses across human patient datasets and multiple experimental models of TTS reveal that dysregulation of phosphorylation-dependent stress signaling constitutes a conserved and unifying molecular feature of the disease. Across species and disease contexts, pathways related to MAPK activation, protein phosphorylation, regulated cell death, and mitochondrial stress were consistently enriched, indicating convergence on stress-responsive signaling networks rather than disparate model-specific alterations. Given this convergence on phosphorylation-dependent pathways, we reasoned that disruption of counter-regulatory phosphatase activity might represent a critical, yet underappreciated, component of the TTS stress response. Consistent with this concept, exposure of cardiomyocytes to plasma from patients with TTS resulted in rapid suppression of PP2A activity without changes in PP2A-C protein abundance. Together, these findings suggest that circulating stress-associated factors in TTS acutely impair PP2A-dependent phosphorylation homeostasis, thereby sensitizing cardiomyocytes to downstream injury pathways.

A defining feature of TTS is its abrupt onset, striking regional heterogeneity, and spontaneous recovery, yet the molecular basis underlying these hallmark characteristics has remained poorly defined.^34–36^ Our findings provide evidence that PP2A activity is dynamically and region-specifically regulated during the course of stress-induced cardiomyopathy. In an established ISO-induced TTS model, PP2A activity was most profoundly suppressed during the acute phase, coinciding temporally with peak cardiac dysfunction and myocardial injury, and subsequently recovered in parallel with functional and structural restoration. Notably, PP2A suppression exhibited marked spatial preference, with the most pronounced reduction observed in the apical myocardium, while basal regions were relatively spared. This regional pattern closely mirrored the distribution of myocardial injury and dysfunction, suggesting that localized loss of PP2A activity may contribute to the characteristic apical vulnerability of the stressed heart. The temporal reversibility and spatial restriction of PP2A inactivation provide a mechanistic framework linking catecholamine stress to the distinctive clinical phenotype of TTS. These observations underscore the acute phase as a critical window during which suppression of PP2A activity lowers the threshold for stress-induced injury in vulnerable myocardial segments. Timely restoration of PP2A activity during this phase may therefore represent an opportunity not only to attenuate acute myocardial injury, but also to favorably influence the trajectory of cardiac recovery. Beyond its association with acute TTS, our study provides direct causal evidence that suppression of PP2A activity actively contributes to disease severity. Using complementary genetic and pharmacological loss-of-function approaches, we demonstrate that cardiomyocyte-specific reduction of PP2A activity markedly exacerbates cardiac dysfunction, myocardial injury, and acute mortality in TTS. Conversely, restoration of PP2A activity through pharmacological activation conferred robust cardioprotection across multiple experimental models of TTS. Activation of PP2A by DT-061 selectively restored methylated PP2A-C levels, improved survival, preserved systolic function, and attenuated myocardial injury following both ISO- and EPI-induced stress. Together, these bidirectional genetic and pharmacological interventions establish PP2A activity as a central determinant of myocardial vulnerability during acute catecholamine stress. findings provide a strong rationale for targeting PP2A-dependent signaling during the early phase of TTS as a strategy to blunt acute injury and facilitate recovery.

Given the growing recognition that stress-responsive signaling pathways contribute to catecholamine-induced myocardial injury,^37,38^ we next sought to define the downstream molecular programs regulated by PP2A during acute TTS. To this end, we performed transcriptomic profiling of cardiac tissues from ISO-induced TTS mice with or without pharmacological activation of PP2A. These analyses revealed robust and concurrent activation of gene programs related to programmed cell death, autophagy, and mitochondrial dysfunction in catecholamine-stressed hearts, indicating engagement of a coordinated stress-response network rather than isolated pathological processes. Importantly, pharmacological activation of PP2A broadly attenuated these transcriptional stress signatures, positioning PP2A as an upstream regulator that constrains multiple converging injury pathways rather than a single downstream effector.

Among regulated cell death modalities, ferroptosis emerged as a dominant feature of acute TTS in our models, characterized by excessive lipid peroxidation and impaired antioxidant defense. This observation is consistent with accumulating evidence implicating iron-dependent lipid oxidative injury as a critical mechanism of acute cardiac damage under ischemic, inflammatory, and catecholaminergic stress conditions.^39–41^ In contrast, canonical markers of apoptosis and pyroptosis were minimally altered, underscoring the specificity of ferroptosis in this context. Notably, restoration of PP2A activity markedly suppressed ferroptosis-associated molecular signatures, identifying PP2A as an endogenous brake on lipid peroxidation-driven cardiomyocyte injury during acute stress. In parallel, our data reveal pronounced activation of ferritinophagy, a selective autophagic pathway governing iron recycling through NCOA4-mediated ferritin degradation.^42–44^ Acute catecholamine stress resulted in degradation of both cytosolic and mitochondrial ferritin pools, increased autophagic flux, and accumulation of labile iron, changes that were exacerbated by PP2A deficiency and attenuated by PP2A activation. These findings suggest that excessive ferritin degradation represents a key downstream mechanism through which PP2A activity regulates myocardial vulnerability in TTS.

Given mitochondrial dysfunction also represents a major downstream and convergent target of PP2A-regulated stress responses in acute TTS and the intimate dependence of mitochondrial redox balance on intracellular iron homeostasis, excessive ferritin degradation and labile iron accumulation are expected to directly sensitize mitochondria to oxidative damage during acute stress.^45, 46^ Previous studies have consistently implicated mitochondrial vulnerability, including impaired bioenergetics, altered mitochondrial dynamics, and oxidative stress, in stress-induced cardiomyopathy and TTS, highlighting mitochondria as a critical determinant of cardiomyocyte resilience under catecholamine stress.^47, 48^ In line with these observations, at the ultrastructural level, acute TTS was characterized by profound mitochondrial abnormalities, including mitochondrial swelling, disruption of the outer mitochondrial membrane, and accumulation of autophagic structures. Notably, such ultrastructural features are increasingly recognized as hallmarks of ferroptosis-associated mitochondrial injury, linking mitochondrial damage directly to iron-dependent lipid peroxidation and oxidative stress. These changes and loss of mitochondria-related proteostasis were markedly alleviated by restoration of PP2A activity, whereas PP2A deficiency further aggravated mitochondrial injury. At the cellular level, suppression of PP2A enhanced cardiomyocyte death, lipid peroxidation and mitochondrial depolarization, whereas restoration of PP2A activity exerted broad cytoprotective effects. In parallel, enhanced ferritinophagy and altered iron handling were consistently observed in vivo and in vitro, accompanied by increased cytosolic and mitochondrial labile iron. Importantly, genetic blockade of ferritinophagy partially rescued cardiomyocyte survival and mitochondrial function, supporting a functional role for iron-dependent pathways downstream of PP2A inactivation.

Collectively, these findings support a model in which PP2A acts as a central signaling integrator that restrains iron-dependent oxidative stress, excessive ferritinophagy, and mitochondrial injury during acute catecholamine exposure. Disruption of PP2A activity removes this protective constraint, permitting coordinated activation of ferroptosis and mitochondrial stress responses that collectively drive cardiomyocyte injury in TTS.

Our study further identifies JNK-MAPK signaling as a critical downstream effector linking PP2A inactivation to cardiomyocyte injury. Transcriptomic analyses highlighted MAPK pathway activation in TTS hearts, prompting targeted investigation of MAPK signaling components. Among these, JNK activation was most prominently induced by catecholamine stress and selectively suppressed by PP2A activation. Pharmacological inhibition of JNK markedly improved cardiac function, reduced myocardial injury, and attenuated ferritinophagy and mitochondria-associated alterations in both pharmacological and genetic models of PP2A inhibition.

These observations are consistent with the established role of JNK as a stress-activated kinase integrating oxidative stress, mitochondrial dysfunction, and cell death pathways.^49,50^ In the context of TTS, our data suggest that PP2A restrains excessive JNK activation, thereby limiting downstream iron-dependent and mitochondrial injury. Thus, these findings indicate that JNK represents a major and therapeutically tractable node within this signaling network.

Although TTS is generally considered a reversible cardiomyopathy, acute complications including heart failure, arrhythmias, and cardiogenic shock remain clinically significant, and no disease-specific therapies are currently available.^51,52^ The identification of PP2A inactivation as a central and reversible molecular event in acute TTS raises the possibility that transient modulation of PP2A-dependent signaling may hold therapeutic relevance during the vulnerable acute phase.

In this regard, our pharmacological studies using DT-061 provide an important proof of concept. DT-061 selectively enhanced PP2A activity without altering PP2A protein abundance and consistently attenuated cardiac dysfunction, ferritinophagy, and mitochondrial dysfunction across multiple experimental models. These effects were observed when PP2A activation was applied in a temporally restricted manner, aligning with the acute and self-limited nature of TTS. While DT-061 is not proposed here as a clinical treatment, these findings support the concept that restoring phosphatase balance during acute catecholamine stress may mitigate myocardial injury and accelerate functional recovery.

Similarly, inhibition of JNK signaling effectively rescued cardiac function and cellular injury in both pharmacological and genetic models of PP2A deficiency, positioning JNK as a downstream effector that may be amenable to therapeutic modulation. Importantly, JNK inhibition attenuated multiple converging pathological processes rather than a single downstream endpoint, highlighting the advantage of targeting signaling hubs within stress-response networks. Together, these observations suggest that transient targeting of the PP2A–JNK axis may represent a rational strategy to blunt acute myocardial injury in TTS, warranting further investigation.

## STUDY LIMITATIONS AND FUTURE DIRECTIONS

Several limitations of the present study should be acknowledged. First, although our data demonstrate coordinated regulation of ferroptosis, ferritinophagy, and mitochondrial dysfunction downstream of PP2A inactivation, the precise causal relationships among these processes remain to be fully delineated. Future studies employing more selective genetic and temporal interventions will be required to dissect the hierarchical organization of iron handling and mitochondrial injury in TTS. Second, our experimental models primarily focus on the acute phase of catecholamine-induced stress cardiomyopathy. While this approach closely mirrors the clinical onset of TTS, longer-term studies will be necessary to determine whether transient modulation of PP2A-dependent signaling influences myocardial recovery or susceptibility to recurrent episodes. In addition, although TTS predominantly affects postmenopausal women, sex-specific mechanisms were not systematically addressed in the current study and represent an important area for future investigation. Finally, while DT-061 and SP600125 served as valuable pharmacological tools to interrogate PP2A-JNK signaling, translation of these findings to clinical practice will require the development of agents with established safety profiles and precise temporal control. Nevertheless, the consistency of our findings across human samples, animal models, and cardiomyocytes supports the relevance of this signaling axis to human disease.

## CONCLUSIONS

In conclusion, our study identifies reversible inactivation of PP2A as a central molecular event in the pathogenesis of acute TTS. Loss of PP2A activity links catecholamine stress to maladaptive activation of JNK-MAPK signaling, ferritinophagy-mediated ferroptosis, and mitochondrial dysfunction in cardiomyocytes. Importantly, restoration of PP2A activity or downstream inhibition of JNK effectively mitigates these converging pathological processes and preserves cardiac function across multiple experimental systems. These findings establish PP2A as a critical integrator of stress-responsive signaling in the heart and provide a unifying mechanistic framework for the reversible myocardial injury characteristic of TTS, highlighting PP2A activation as a promising target for transient therapeutic modulation during the acute phase of the disease.

## ACKNOWLEDGMENTS

The authors thank Dr. Puja K. Mehta for providing human samples and Dr. William M. Chilian for generously sharing the RNA-seq dataset from the TAC model.

