## Supplementary Figures for "Protein Phosphatase 2A Activation Attenuates Acute Myocardial Injury in Takotsubo Syndrome by Modulating Ferroptosis and Mitochondrial Injury in Cardiomyocytes"

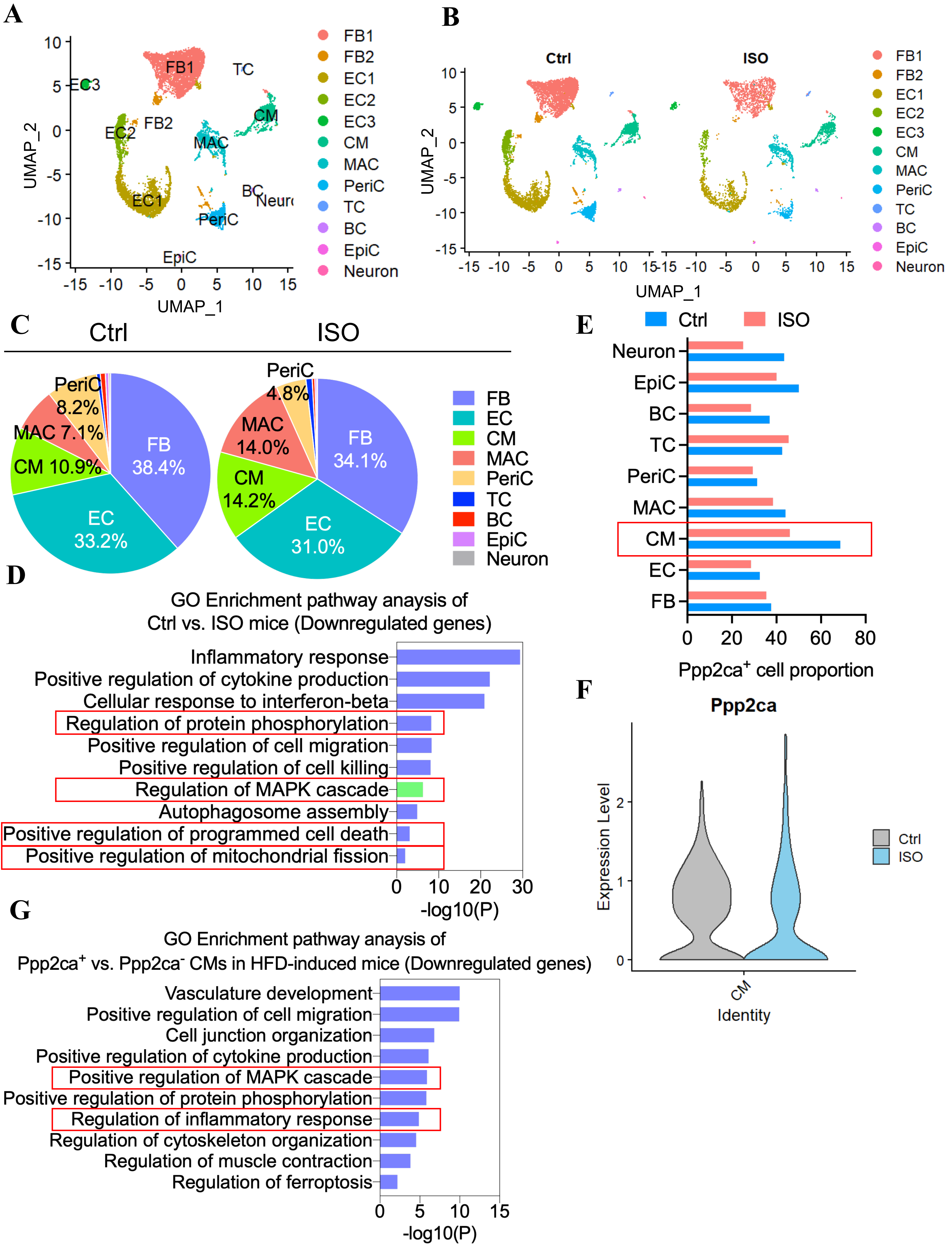


**Supplementary Figure 1. Single-cell transcriptomic analysis reveals stress-induced phosphorylation-related signaling alterations in ISO-treated hearts. A,** UMAP visualization of single-cell RNA-sequencing data from control and ISO-treated mouse hearts. **B,** Split UMAP visualization comparing control and ISO-treated hearts. **C,** Relative proportions of major cardiac cell populations in control and ISO-treated hearts. **D,** GO enrichment analysis of downregulated genes in ISO-treated hearts. E, GO enrichment analysis of differentially expressed genes in cardiomyocytes from ISO-treated hearts, showing enrichment of stress-responsive and phosphorylation-related biological processes. E, Proportion of Ppp2ca-positive cells across major cardiac cell populations in control and ISO-treated hearts. **F,** Violin plot showing Ppp2ca expression levels in cardiomyocytes from control and ISO-treated hearts. **G,** GO enrichment analysis of downregulated genes in Ppp2ca-positive versus Ppp2ca-negative cardiomyocytes from ISO-treated hearts. CM, cardiomyocytes; EC, endothelial cells; FB, fibroblasts; MAC, macrophages; PeriC, pericytes; TC, T cells; BC, B cells; EpiC, epicardial cells; ISO, isoprenaline; UMAP, uniform manifold approximation and projection; GO, Gene Ontology.

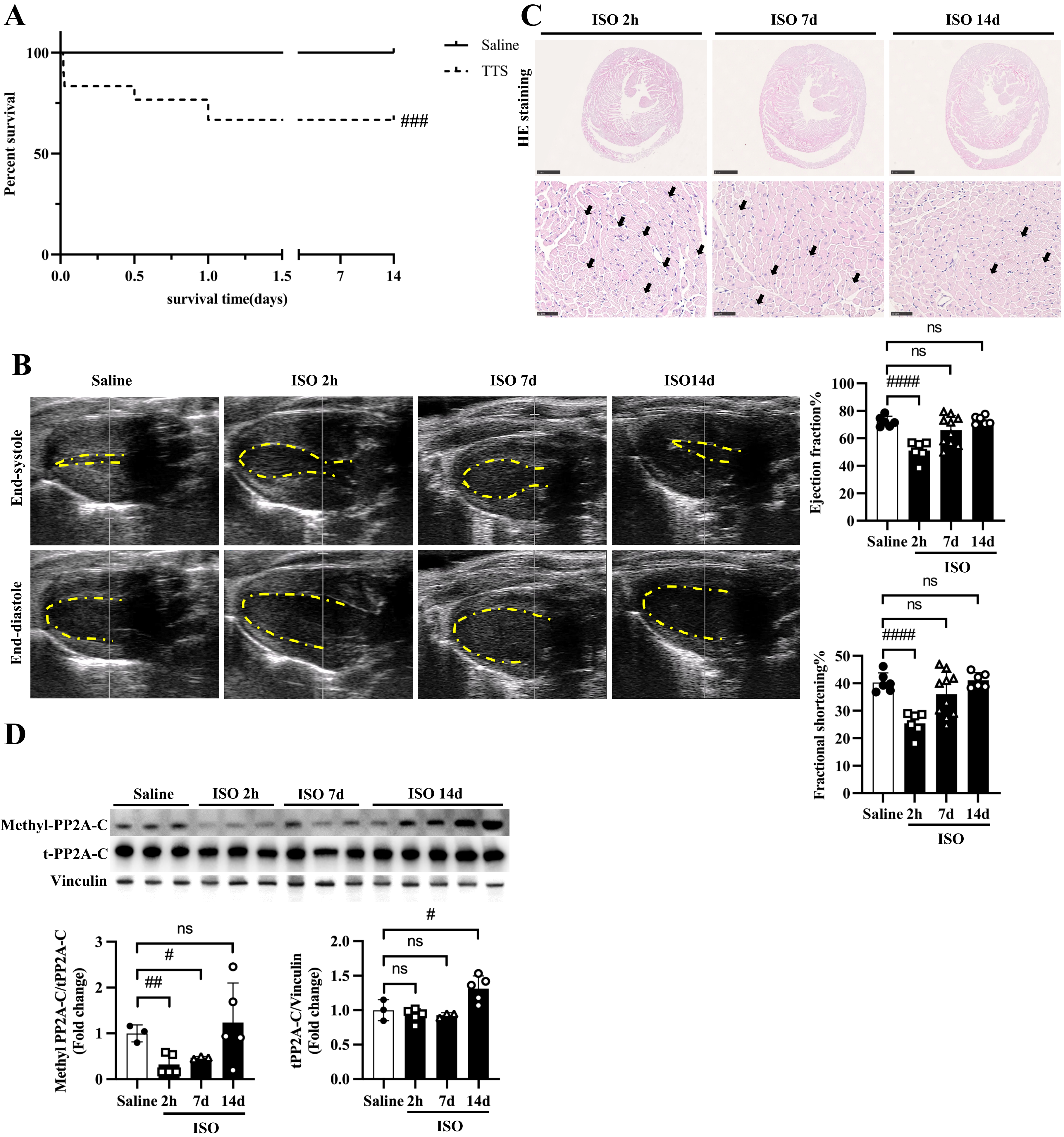


**Supplementary Figure 2. Temporal recovery of cardiac function, myocardial structure, and PP2A activity following ISO-induced TTS. A,** Kaplan–Meier survival curve showing mortality over time following ISO administration. **B,** Echocardiographic assessment of cardiac function at 2 hours, 7 days, and 14 days after ISO treatment, including quantification of EF and FS. **C,** HE staining of left ventricular sections at 2 hours, 7 days, and 14 days after ISO treatment, demonstrating progressive recovery of myocardial structure over time. **D,** Immunoblot analysis of methyl-PP2A-C and total PP2A-C in cardiac tissue at 2 hours, 7 days, and 14 days after ISO treatment, with corresponding quantification. Data are presented as mean ± SEM. #*P*<0.05; ##*P*<0.01; ###*P*<0.001; ####*P*<0.0001; ns, not significant.

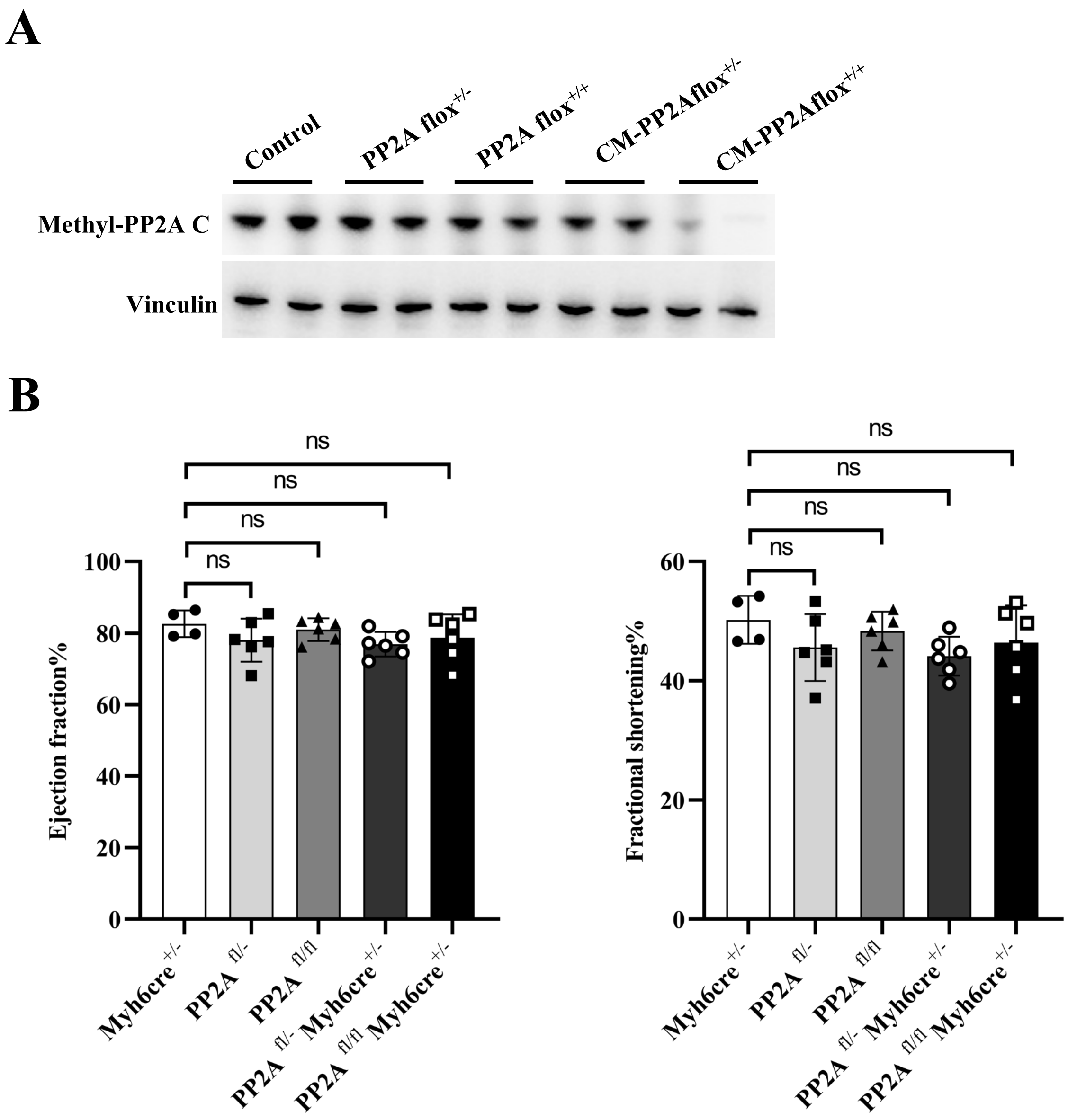


**Supplementary Figure 3. Baseline cardiac phenotype and functional validation of PP2A suppression models. A,** Immunoblot analysis of total PP2A-C (t-PP2A-C) in cardiac tissue from control (Myh6-Cre; $PP2Afl^{+/-}$; $PP2Afl^{+/+}$) and cardiomyocyte-specific PP2A-Cα–deficient mice (CM-$PP2Afl^{+/-}$; CM-$PP2Afl^{+/-}$). **B,** Echocardiographic assessment of cardiac function control (Myh6-Cre; $PP2Afl^{+/-}$; $PP2Afl^{+/+}$) and cardiomyocyte-specific PP2A-Cα–deficient mice (CM-$PP2Afl^{+/-}$; CM-$PP2Afl^{+/-}$) under basal conditions following tamoxifen administration, including EF and FS. Data are presented as mean±SEM. ns, not significant.

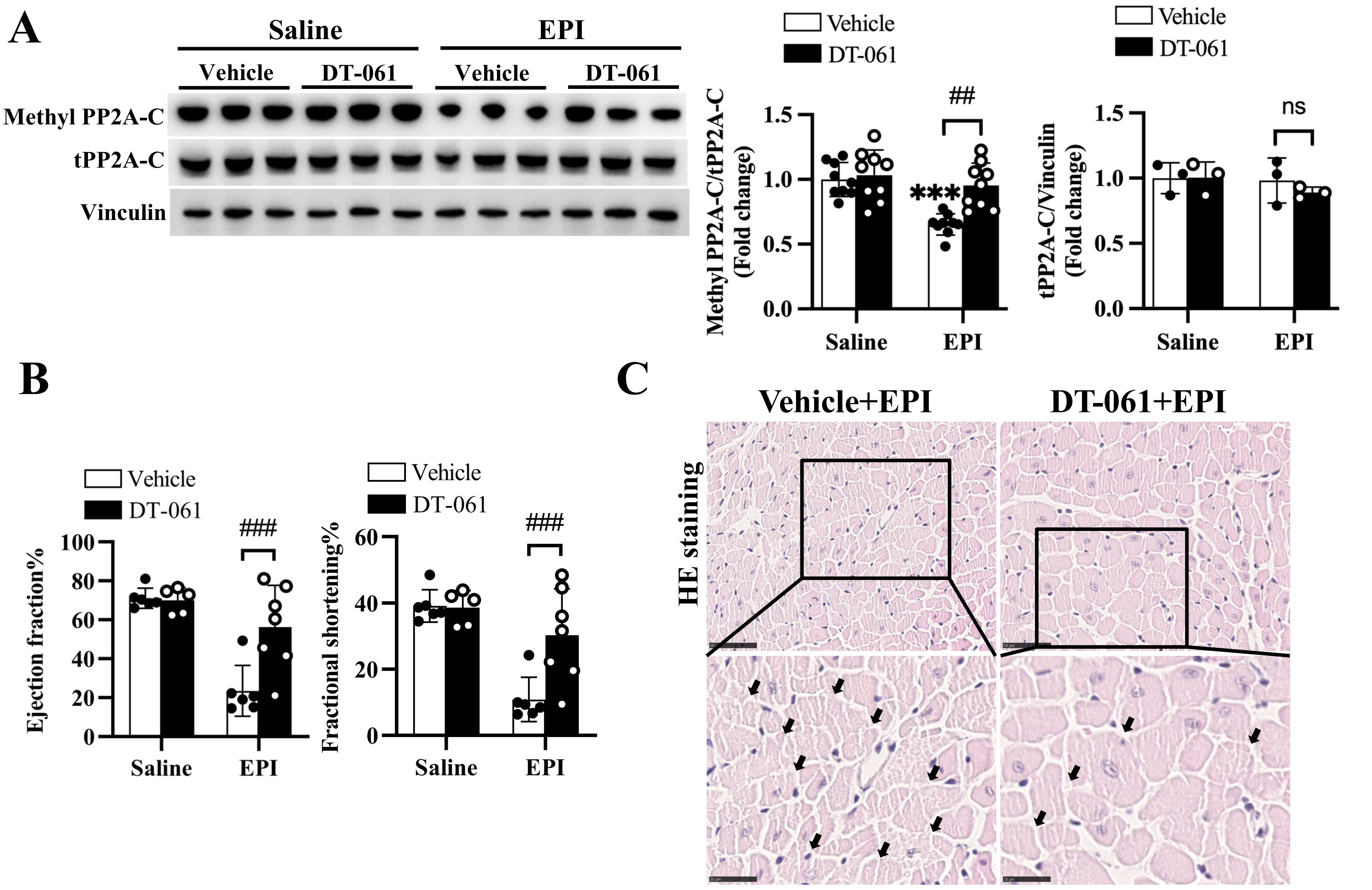


**Supplementary Figure 4. PP2A activation confers cardioprotection in an epinephrine-induced TTS model. A,** Immunoblot analysis of methyl-PP2A-C and total PP2A-C in cardiac tissue from EPI-induced TTS followed by treatment with DT-061, with corresponding quantification normalized to vinculin. **B,** Echocardiographic assessment of cardiac function in EPI-induced TTS mice followed by treatment with DT-061, including quantification of EF and FS. **C,** HE staining of cardiac sections from the indicated groups. Arrows indicate areas of myocardial injury. Data are presented as mean±SEM. *$P$<0.05; **$P$<0.01; ***$P$<0.001; ****$P$<0.0001; #$P$<0.05; ##$P$<0.01; ###$P$<0.001; ####$P$<0.0001; ns, not significant.

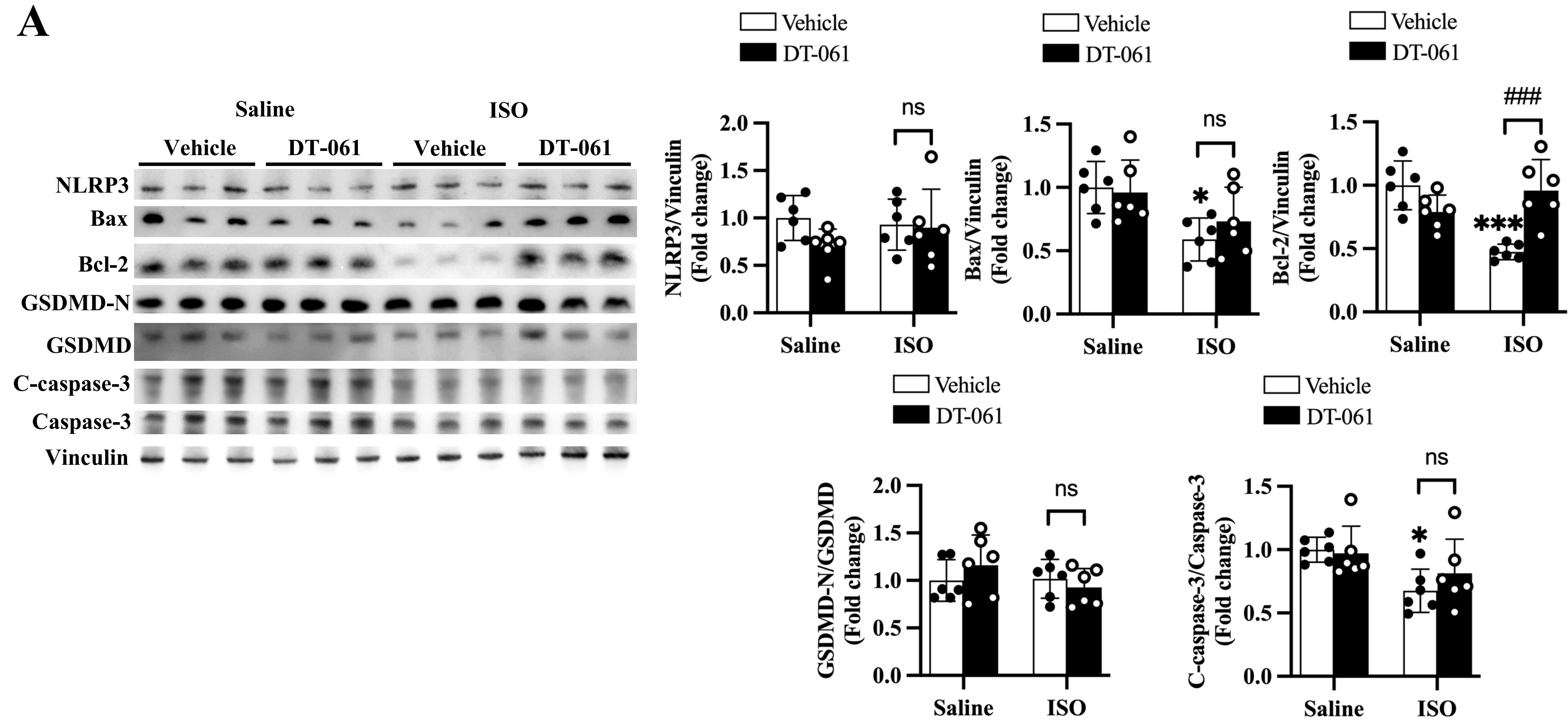


**Supplementary Figure 5 (related to Figure 5). Apoptosis and pyroptosis are not major contributors to acute TTS. A,** Immunoblot analysis of apoptosis-related and pyroptosis-related markers in cardiac tissue from saline- and ISO-treated mice followed by DT-061 admnistration. Data are presented as mean±SEM. Data are presented as mean±SEM. *$P$<0.05; **$P$<0.01; ***$P$<0.001; ****$P$<0.0001; #$P$<0.05; ##$P$<0.01; ###$P$<0.001; ####$P$<0.0001; ns, not significant.

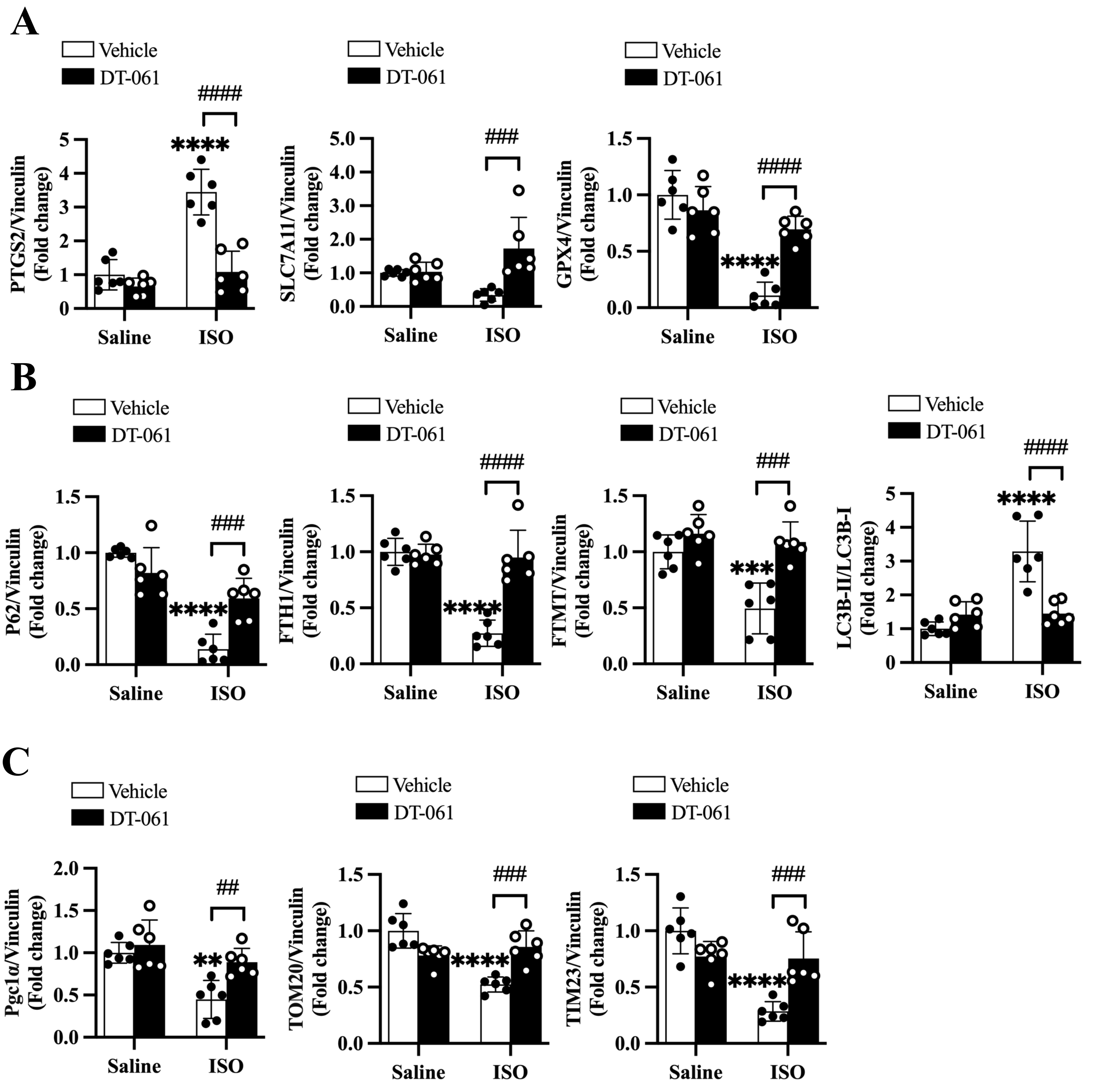


**Supplementary Figure 6 (related to Figure 5). PP2A activation attenuates ferroptosis, ferritinophagy, and mitochondrial dysfunction. A,** Quantification of ferroptosis-related proteins corresponding to Figure 5F. **B,** Quantification of ferritinophagy-related proteins corresponding to Figure 5G. **C,** Quantification of mitochondrial proteins corresponding to Figure 5J. Data are presented as mean±SEM. **P*<0.05; ***P*<0.01; ****P*<0.001; *****P*<0.0001; #*P*<0.05; ##*P*<0.01; ###*P*<0.001; ####*P*<0.0001; ns, not significant.

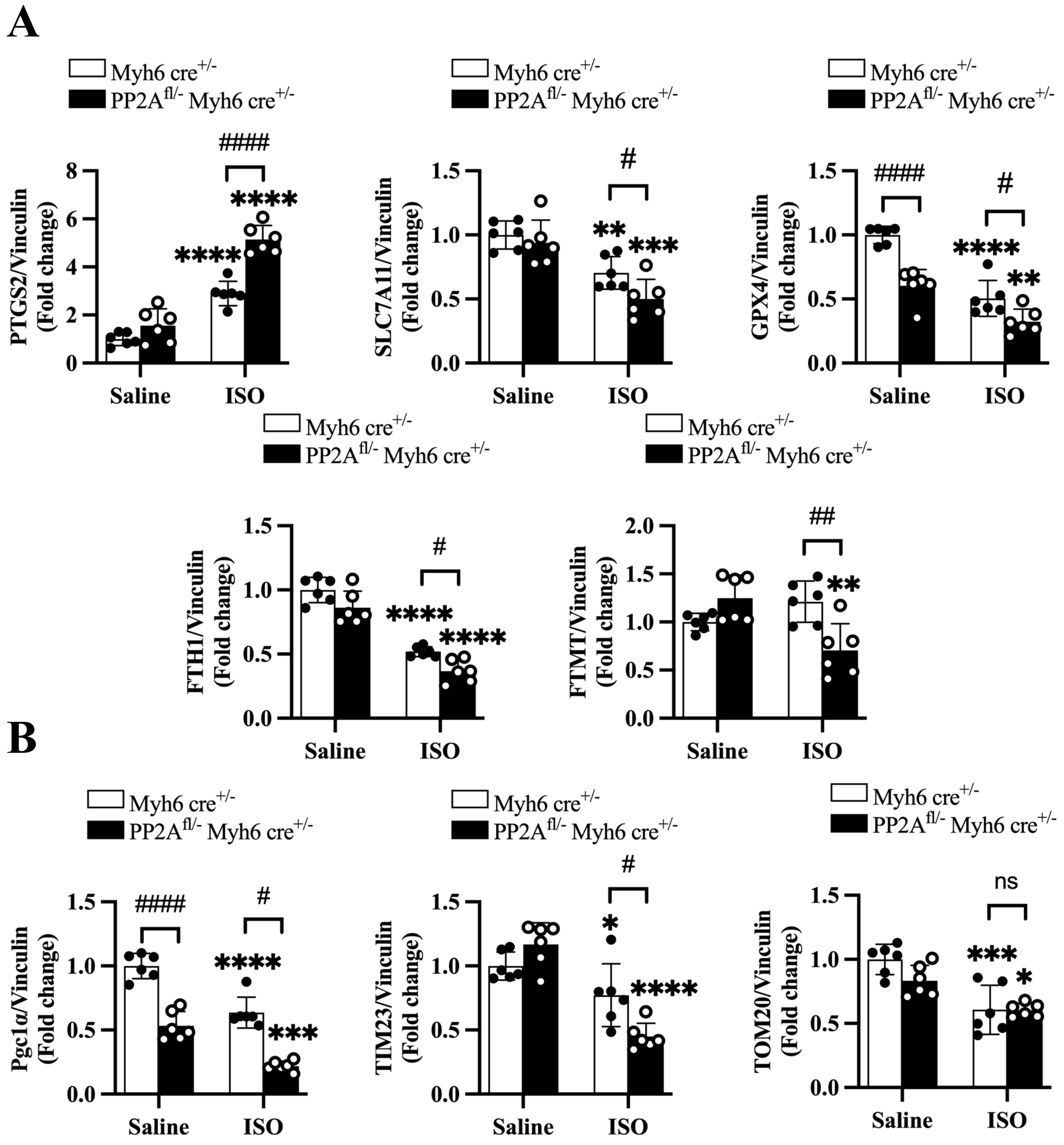


**Supplementary Figure 7 (related to Figure 5). Genetic suppression of PP2A exacerbates ferritinophagy and mitochondrial injury. A,** Quantification of ferritinophagy-related proteins corresponding to Figure 5H. **B,** Quantification of mitochondrial proteins corresponding to Figure 5K. Data are presented as mean±SEM. **P*<0.05; ***P*<0.01; ****P*<0.001; *****P*<0.0001; #*P*<0.05; ##*P*<0.01; ###*P*<0.001; ####*P*<0.0001; ns, not significant.

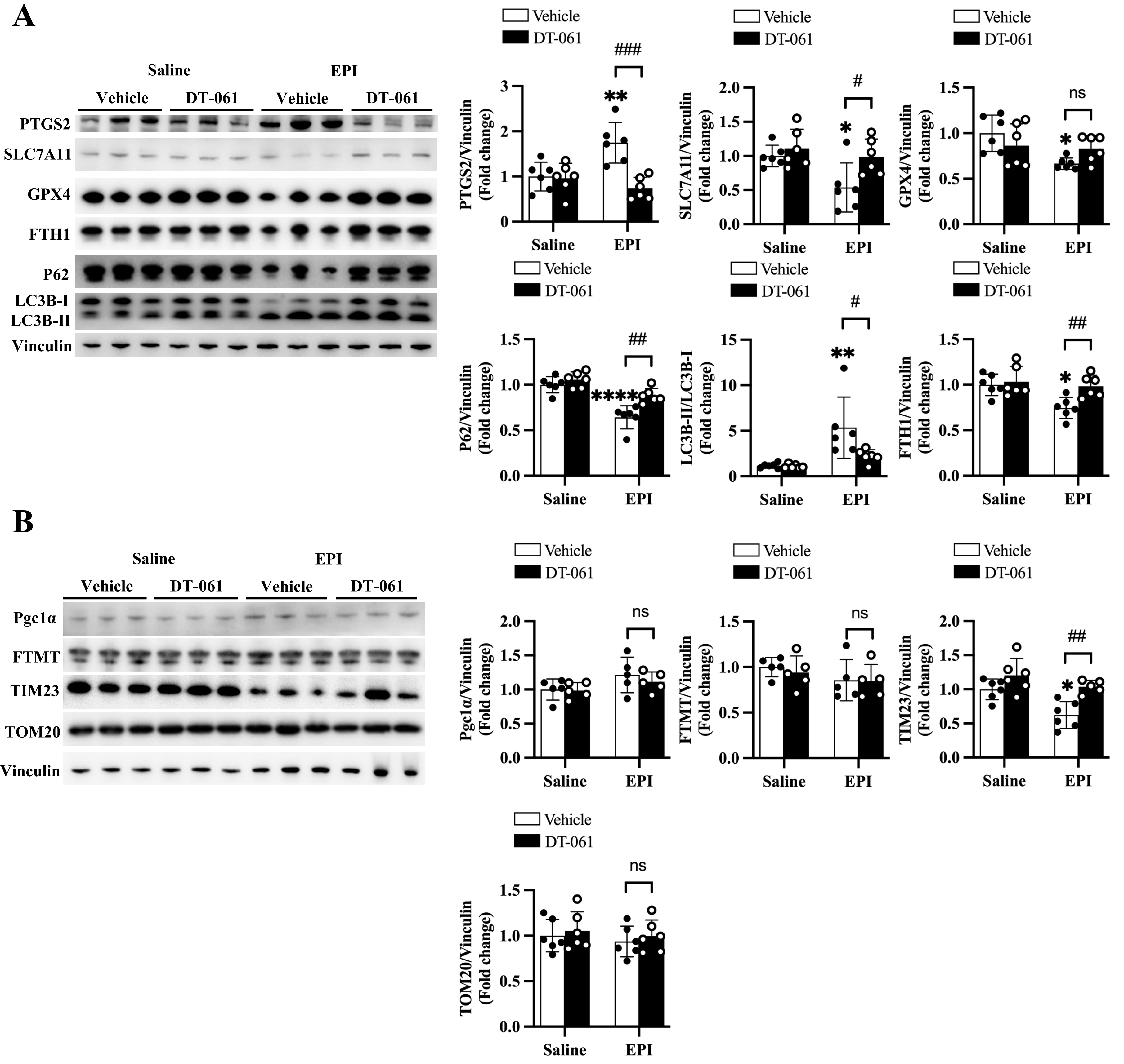


**Supplementary Figure 8 (related to Figure 5). PP2A-dependent regulation of ferroptosis and mitochondrial injury in an epinephrine-induced TTS model. A,** Immunoblot analysis of ferroptosis and ferritinophagy-related proteins in cardiac tissue from DT-061-treated EPI-induced mice. **B,** Immunoblot analysis of mitochondrial proteins in cardiac tissue from DT-061-treated EPI-induced mice. Data are presented as mean±SEM. **P*<0.05; ***P*<0.01; ****P*<0.001; *****P*<0.0001; #*P*<0.05; ##*P*<0.01; ###*P*<0.001; ####*P*<0.0001; ns, not significant.

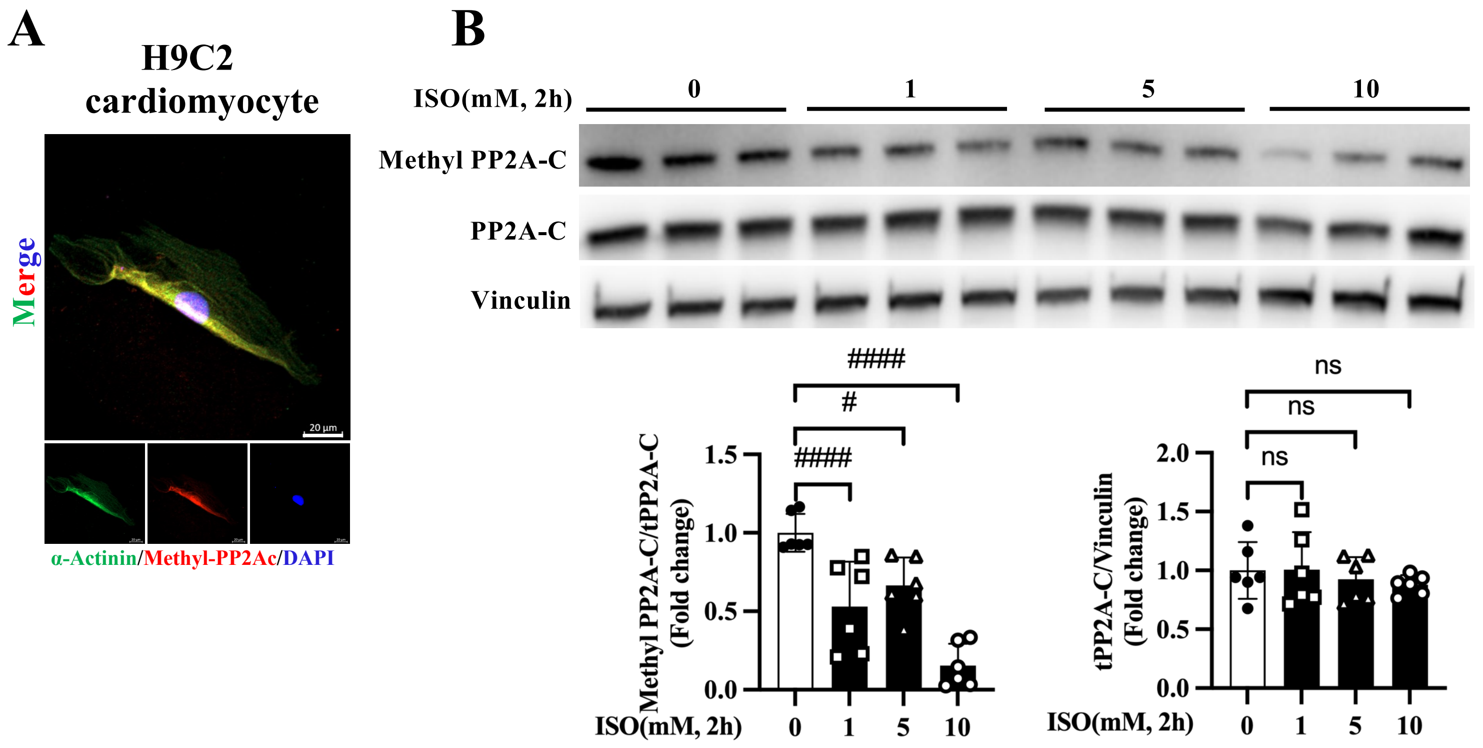


**Supplementary Figure 9 (related to Figure 6). Establishment of an in vitro model of PP2A regulation under catecholamine stress. A,** Immunofluorescence staining of methyl-PP2A-C in H9C2 cells. **B,** Immunoblot analysis of methyl-PP2A-C in H9C2 cells treated with increasing concentrations of ISO. Data are presented as mean±SEM. #*P*<0.05; ##*P*<0.01; ###*P*<0.001; ####*P*<0.0001; ns, not significant.

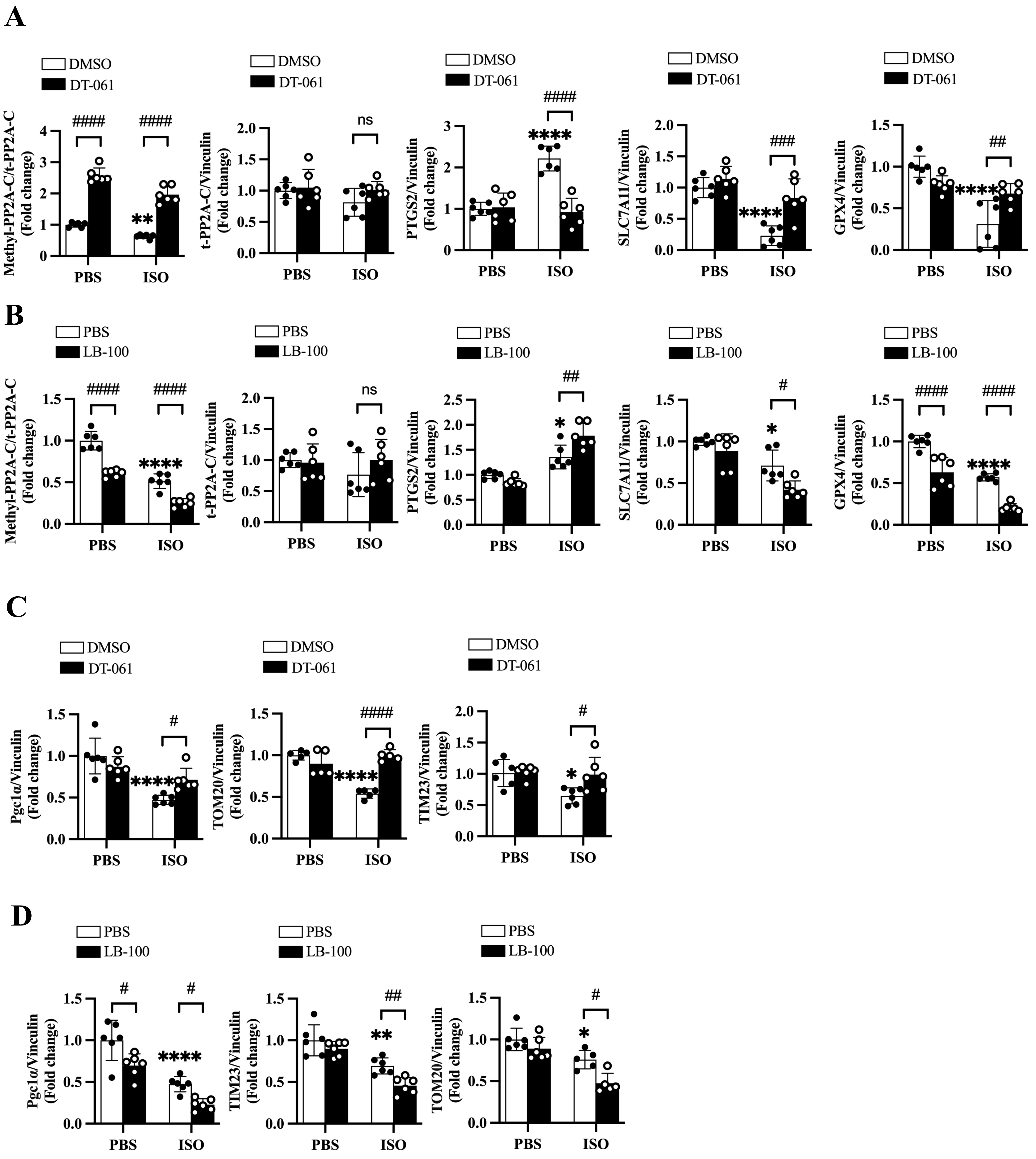


**Supplementary Figure 10 (related to Figure 6). PP2A activity modulates ferroptosis and mitochondrial signaling in H9C2 cells. A,** Quantification of ferroptosis-related proteins corresponding to Figure 6C. **B,** Quantification of ferroptosis-related proteins corresponding to Figure 6D. **C,** Quantification of mitochondrial proteins corresponding to Figure 6G. **D,** Quantification of mitochondrial proteins corresponding to Figure 6H. Data are presented as mean±SEM. **P*<0.05; ***P*<0.01; ****P*<0.001; *****P*<0.0001; #*P*<0.05; ##*P*<0.01; ###*P*<0.001; ####*P*<0.0001; ns, not significant.

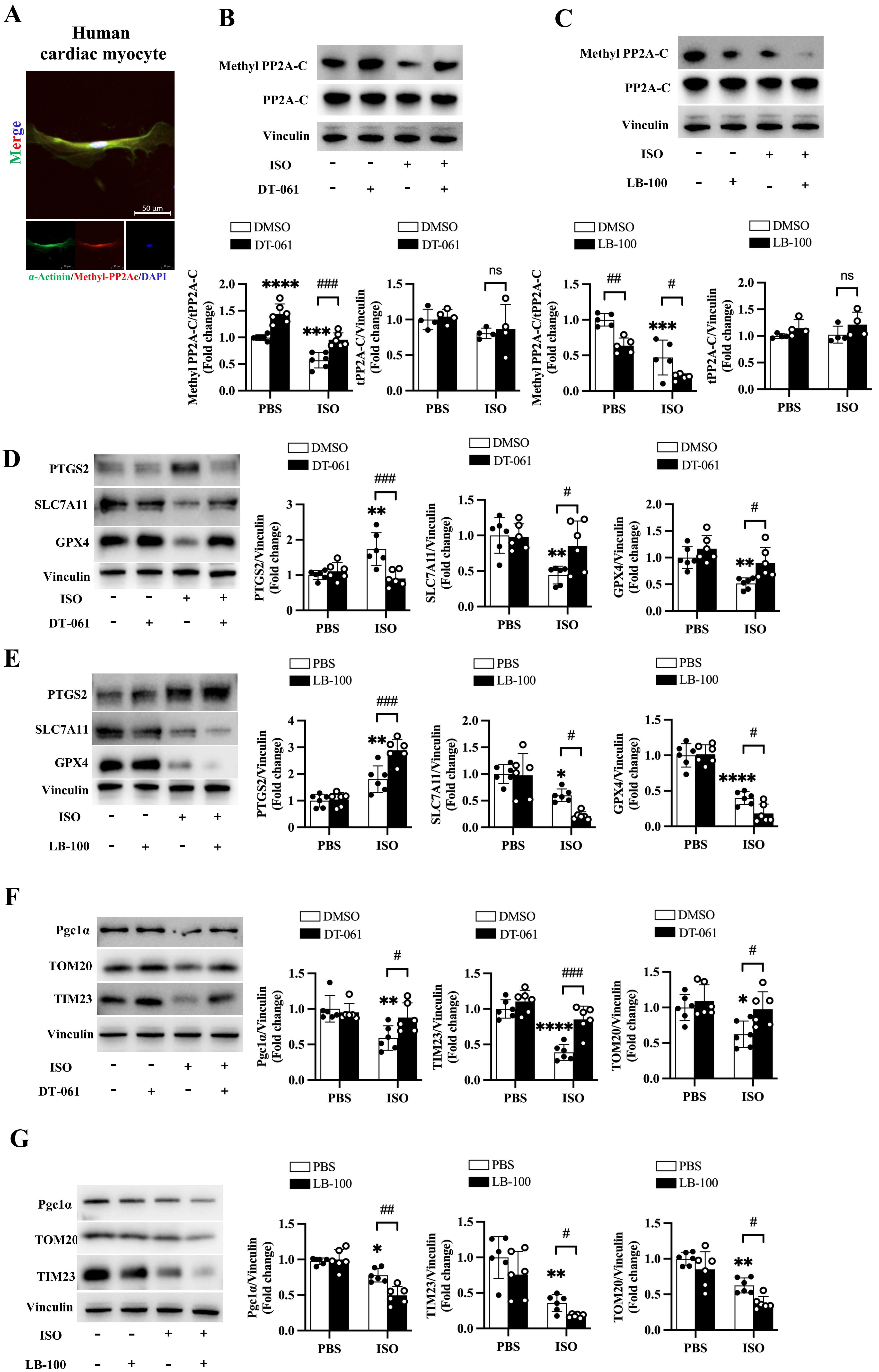
A
Human cardiac myocyte
Merge
50 µm
α-Actinin/Methyl-PP2Ac/DAPI
B
Methyl PP2A-C
PP2A-C
Vinculin
ISO
DT-061
C
Methyl PP2A-C
PP2A-C
Vinculin
ISO
LB-100
DMSO
DT-061
LB-100
Methyl PP2A-C/tPP2A-C (Fold change)
tPP2A-C/Vinculin (Fold change)
****
***
###
##
#
ns
PBS
ISO
D
PTGS2
SLC7A11
GPX4
Vinculin
ISO
DT-061
PTGS2/Vinculin (Fold change)
SLC7A11/Vinculin (Fold change)
GPX4/Vinculin (Fold change)
E
PBS
LB-100
F
Pgc1α
TOM20
TIM23
Vinculin
Pgc1α/Vinculin (Fold change)
TIM23/Vinculin (Fold change)
TOM20/Vinculin (Fold change)
G

**Supplementary Figure 11 (related to Figure 6). Conserved role of PP2A in regulating ferroptosis and mitochondrial dysfunction in human cardiomyocytes. A,** Immunofluorescence staining of methyl-PP2A-C in primary human cardiomyocytes. **B,** Immunoblot analysis of methyl-PP2A-C in primary human cardiomyocytes treated with ISO in the presence or absence of DT-061. **C,** Immunoblot analysis of methyl-PP2A-C and ferroptosis-related proteins in primary human cardiomyocytes treated with LB-100. **D,** Immunoblot analysis of ferroptosis-related protein proteins in primary human cardiomyocytes treated with DT-061, with quantification. **E,** Immunoblot analysis of ferroptosis-related protein proteins in primary human cardiomyocytes treated with LB-100, with quantification. **F,** Immunoblot analysis of mitochondrial proteins in primary human cardiomyocytes treated with DT-061, with quantification. **G,** Immunoblot analysis of mitochondrial proteins in primary human cardiomyocytes treated with LB-100, with quantification. Data are presented as mean±SEM. *P<0.05; **P<0.01; ***P<0.001; ****P<0.0001; #P<0.05; ##P<0.01; ###P<0.001; ####P<0.0001; ns, not significant.

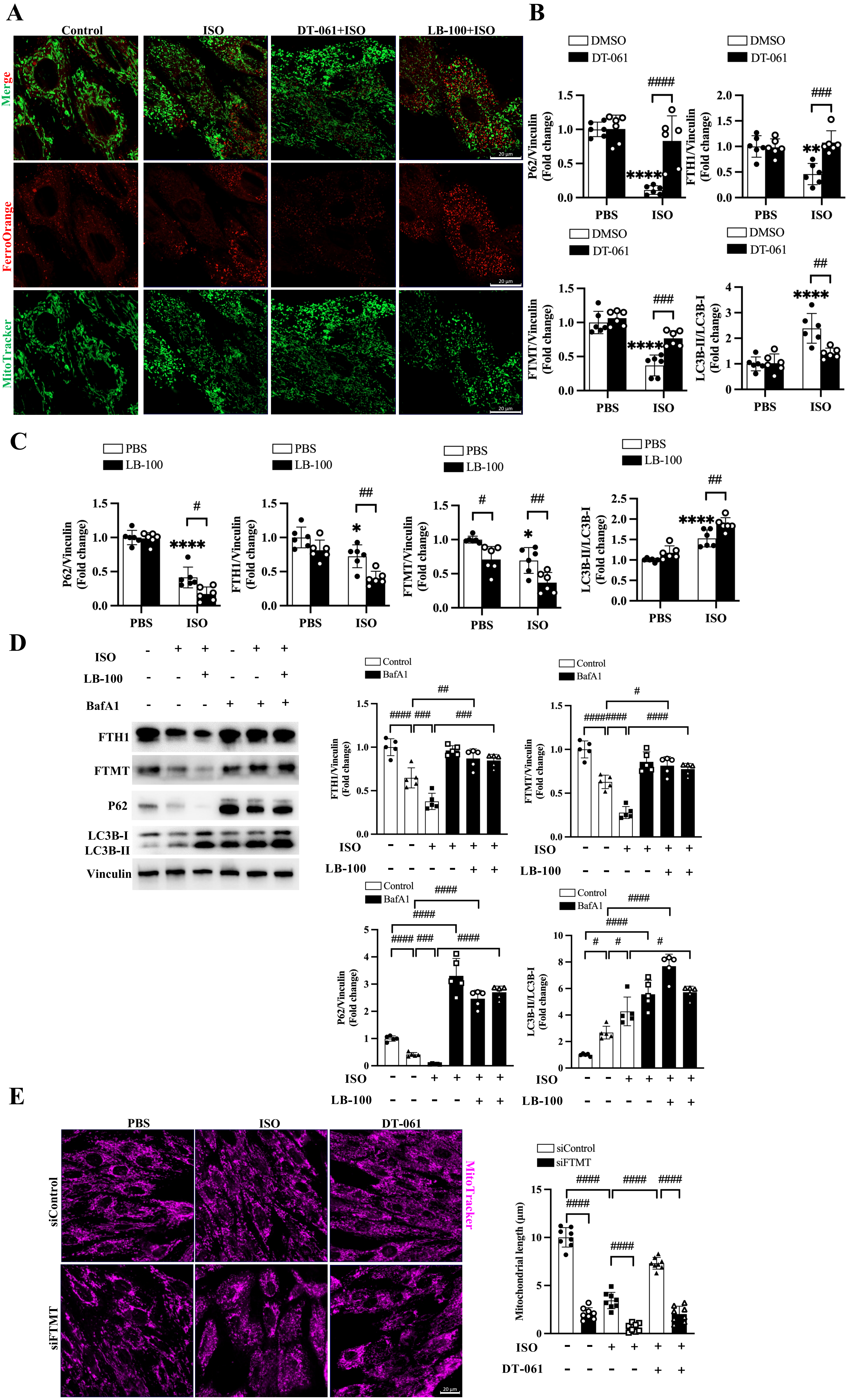

A
Control
ISO
DT-061+ISO
LB-100+ISO
Merge
FerroOrange
MitoTracker
20 μm
B
DMSO
DT-061
PBS
ISO
P62/Vinculin (Fold change)
FTH1/Vinculin (Fold change)
FTMT/Vinculin (Fold change)
LC3B-II/LC3B-I (Fold change)
C
PBS
LB-100
D
ISO
LB-100
BafA1
FTH1
FTMT
P62
LC3B-I
LC3B-II
Vinculin
Control
BafA1
E
PBS
ISO
DT-061
siControl
siFTMT
MitoTracker
siControl
siFTMT
Mitochondrial length (μm)
ISO
DT-061

**Supplementary Figure 12 (related to Figure 7). Ferritinophagy mediates PP2A-dependent regulation of iron handling and mitochondrial function in H9C2 cells. A,** Representative images of FerroOrange and MitoTracker co-staining in H9C2 cells. **B,** Immunoblot analysis of ferritinophagy-related proteins in H9C2 cells treated with ISO and DT-061. **C,** Immunoblot analysis of ferritinophagy-related proteins in H9C2 cells treated with ISO and LB-100. **D,** Immunoblot analysis of autophagic flux markers (LC3-II and p62) under LB-100 and ISO administration pretreated with bafilomycin A1 treatment. **E,** Representative images and quantification of of mitochondrial morphology in H9C2 cells following FTMT knockdown. Data are presented as mean±SEM. *P<0.05; **P<0.01; ***P<0.001; ****P<0.0001; #P<0.05; ##P<0.01; ###P<0.001; ####P<0.0001; ns, not significant.

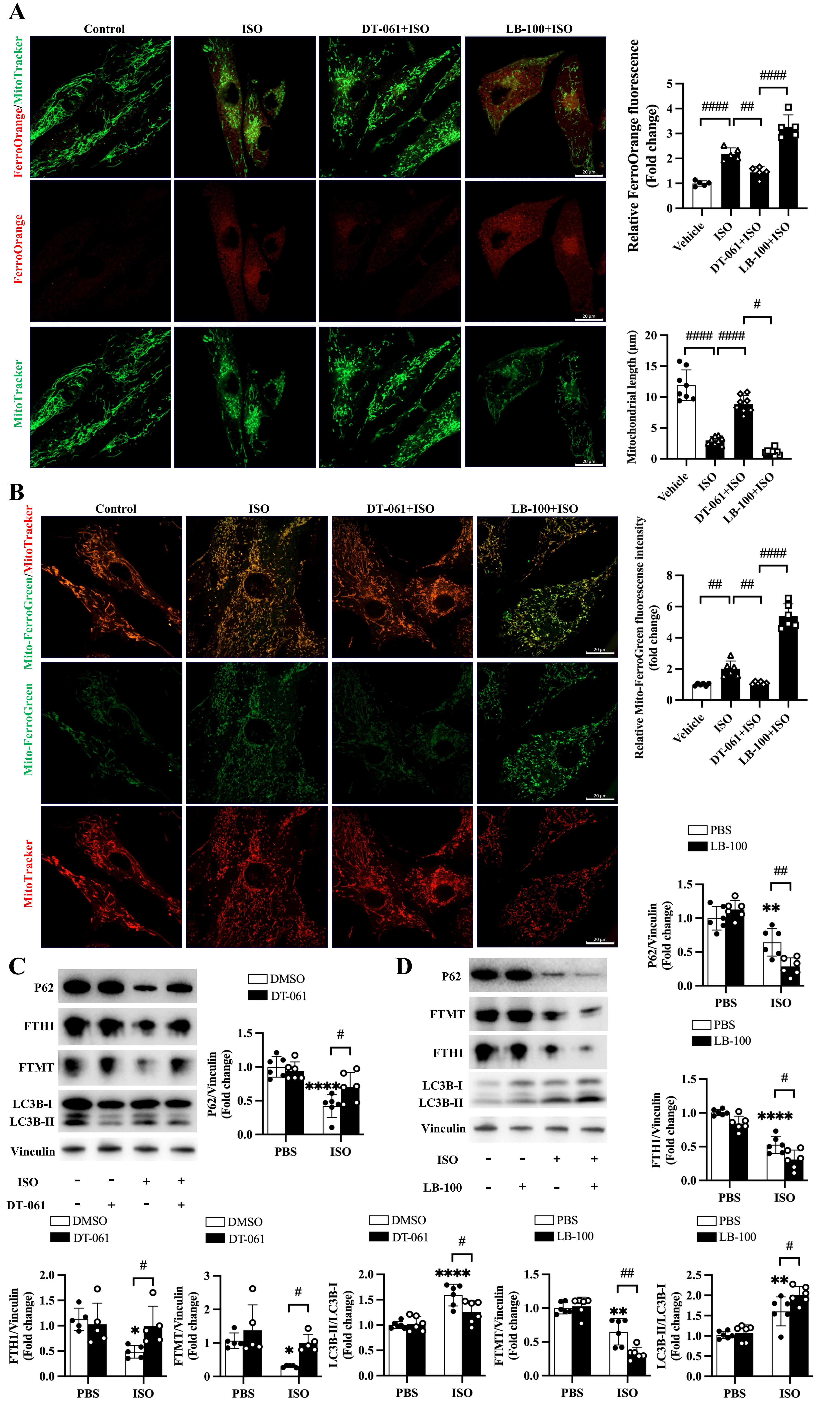

A
Control
ISO
DT-061+ISO
LB-100+ISO
FerroOrange/MitoTracker
FerroOrange
MitoTracker
20 μm
Relative FerroOrange fluorescence (Fold change)
Vehicle
ISO
DT-061+ISO
LB-100+ISO
####
##
####
Mitochondrial length (μm)
####
####
#
B
Control
ISO
DT-061+ISO
LB-100+ISO
Mito-FerroGreen/MitoTracker
Mito-FerroGreen
MitoTracker
Relative Mito-FerroGreen fluorescense intensity (fold change)
##
##
####
PBS
LB-100
P62/Vinculin (Fold change)
**
##
C
P62
FTH1
FTMT
LC3B-I
LC3B-II
Vinculin
ISO
DT-061
DMSO
DT-061
P62/Vinculin (Fold change)
****
#
PBS
ISO
D
P62
FTMT
FTH1
LC3B-I
LC3B-II
Vinculin
ISO
LB-100
FTH1/Vinculin (Fold change)
****
#
FTH1/Vinculin (Fold change)
*
#
FTMT/Vinculin (Fold change)
*
#
LC3B-II/LC3B-I (Fold change)
****
#
FTMT/Vinculin (Fold change)
**
##
LC3B-II/LC3B-I (Fold change)
**
#

**Supplementary Figure 13 (related to Figure 7). PP2A-dependent regulation of iron homeostasis and mitochondrial integrity is conserved in human cardiomyocytes. A,** FerroOrange and MitoTracker co-staining in primary human cardiomyocytes, with quantification. **B,** Representative images and quantification of mitochondrial $Fe^{2+}$ (Mito-FerroGreen) and mitochondrial morphology in human cardiomyocytes under the indicated conditions. **C,** Immunoblot analysis and quantification of ferritinophagy-related proteins in human cardiomyocytes treated with ISO and DT-061. **D,** Immunoblot analysis and quantification of ferritinophagy-related proteins in human cardiomyocytes treated with ISO and LB-100. Data are presented as mean±SEM. *P<0.05; **P<0.01; ***P<0.001; ****P<0.0001; #P<0.05; ##P<0.01; ###P<0.001; ####P<0.0001; ns, not significant.

A

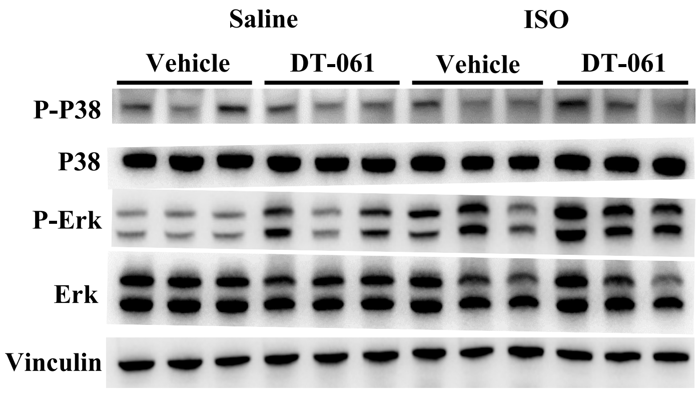


**Supplementary Figure 14 (related to Figure 8). Selective activation of JNK signaling in ISO-induced TTS. A,** Immunoblot analysis of MAPK signaling pathways (p-JNK, p-ERK, p-p38) in TTS mouse hearts pretreated with DT-061 administration.

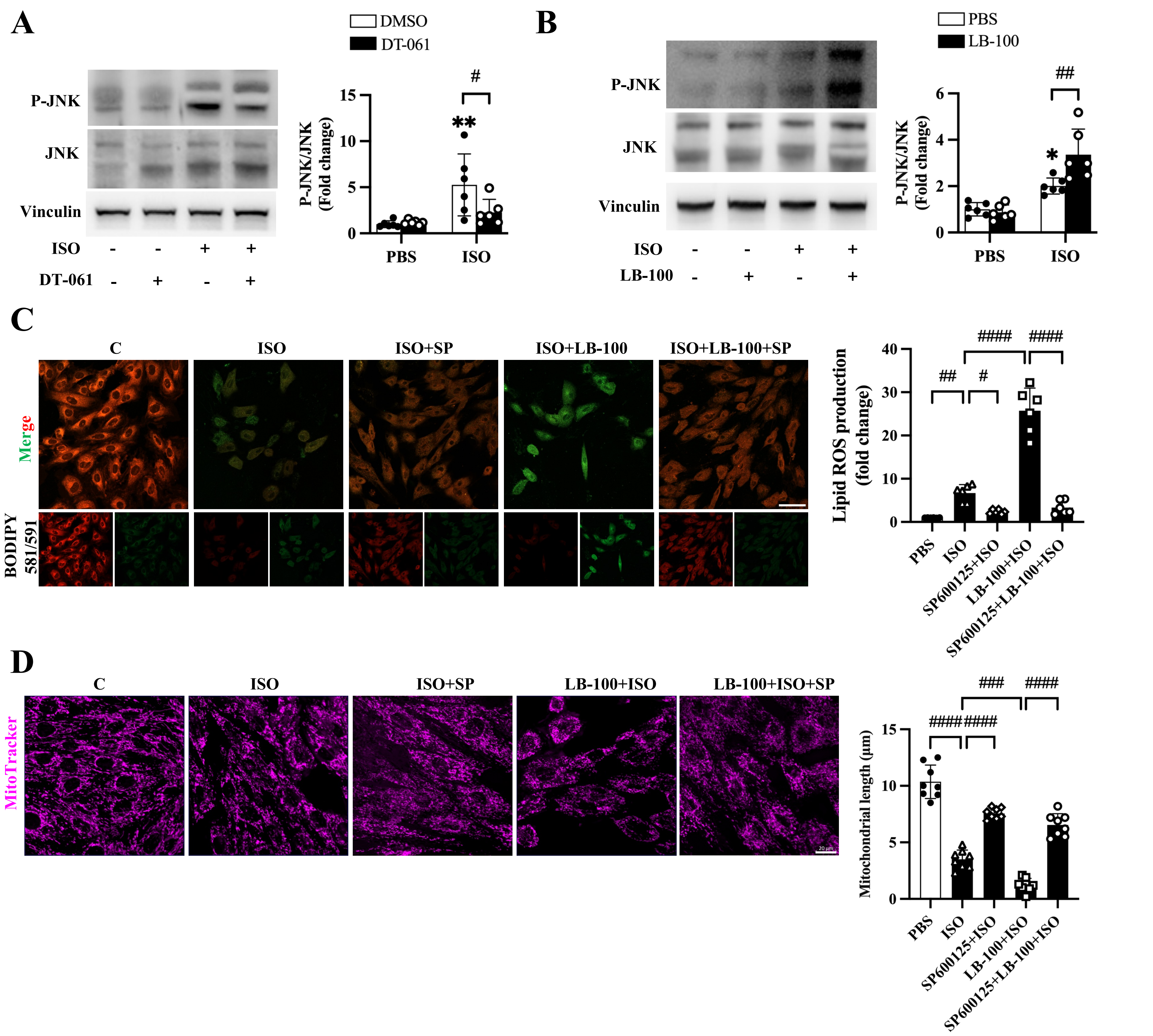


**Supplementary Figure 15 (related to Figure 8). JNK signaling mediates PP2A-dependent cardiomyocyte injury in vitro. A,** Immunoblot analysis of JNK phosphorylation in H9C2 cells following ISO treatment with or without DT-061. **B,** Immunoblot analysis of JNK phosphorylation in H9C2 cells following ISO treatment with or without LB-100. **C,** Representative images and quantification of lipid ROS levels in H9C2 cells treated with ISO and LB-100 in the presence or absence of SP600125. **D,** Assessment of mitochondrial morphology (MitoTracker) following JNK inhibition in H9C2 cells treated with ISO and LB-100 in the presence or absence of SP600125.

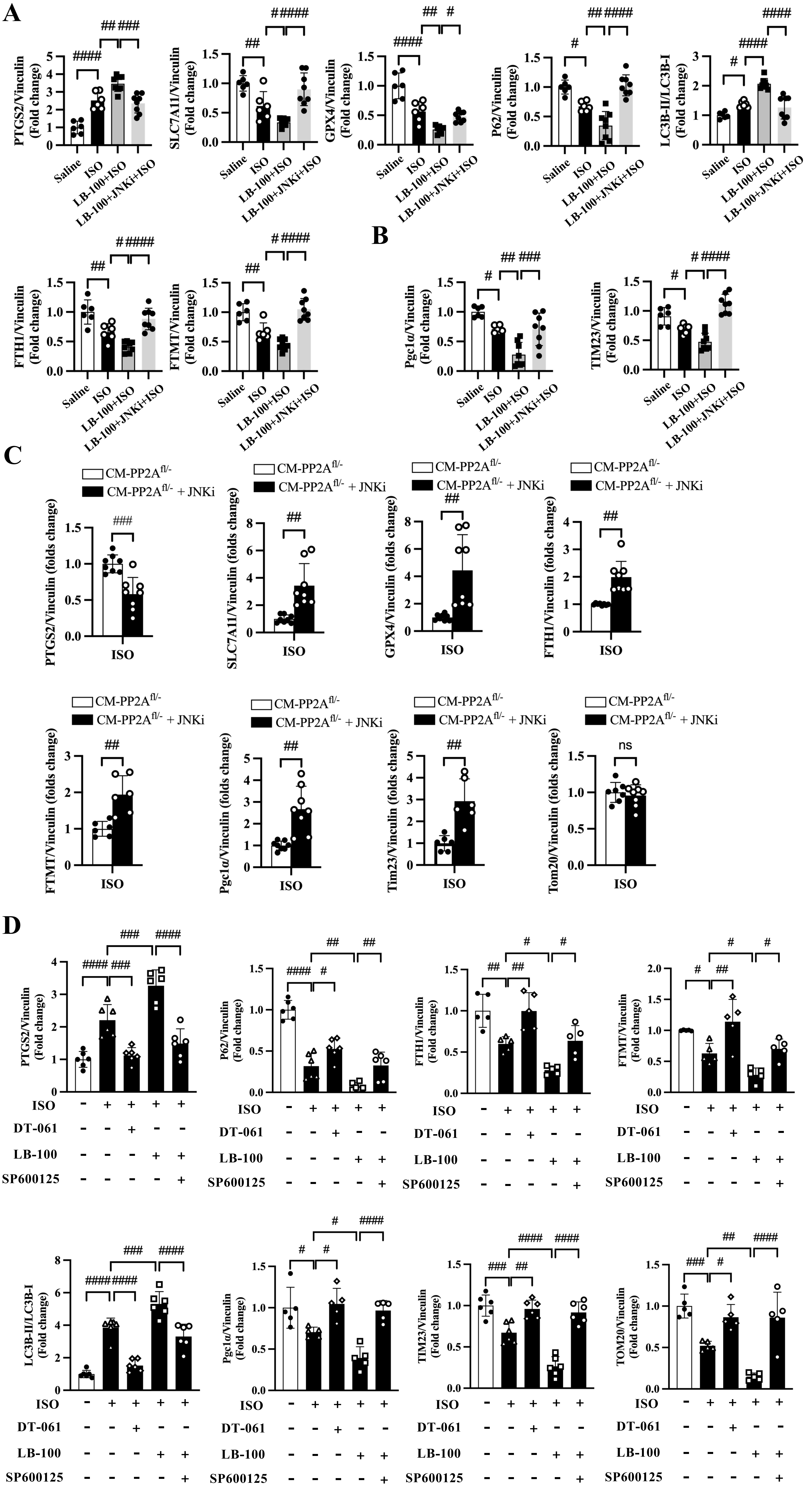
A
PTGS2/Vinculin (Fold change)
SLC7A11/Vinculin (Fold change)
GPX4/Vinculin (Fold change)
P62/Vinculin (Fold change)
LC3B-II/LC3B-I (Fold change)
FTH1/Vinculin (Fold change)
FTMT/Vinculin (Fold change)
Saline
ISO
LB-100+ISO
LB-100+JNKi+ISO
B
Pgc1α/Vinculin (Fold change)
TIM23/Vinculin (Fold change)
C
CM-PP2A fl/-
CM-PP2A fl/- + JNKi
PTGS2/Vinculin (folds change)
SLC7A11/Vinculin (folds change)
GPX4/Vinculin (folds change)
FTH1/Vinculin (folds change)
FTMT/Vinculin (folds change)
Pgc1α/Vinculin (folds change)
Tim23/Vinculin (folds change)
Tom20/Vinculin (folds change)
ns
D
PTGS2/Vinculin (Fold change)
P62/Vinculin (Fold change)
FTH1/Vinculin (Fold change)
FTMT/Vinculin (Fold change)
LC3B-II/LC3B-I (Fold change)
Pgc1α/Vinculin (Fold change)
TIM23/Vinculin (Fold change)
TOM20/Vinculin (Fold change)
ISO
DT-061
LB-100
SP600125

**Supplementary Figure 16 (related to Figure 8). JNK inhibition rescues ferritinophagy and mitochondrial dysregulation induced by PP2A suppression. A,** Quantification of ferritinophagy- and ferroptosis-related proteins in mouse hearts following JNK inhibition under ISO and LB-100 treatment. **B,** Quantification of mitochondrial regulatory proteins following JNK inhibition. **C,** Quantification of ferritinophagy- and mitochondrial-related proteins in PP2A-Cα–deficient mice following JNK inhibition. **D,** Quantification of ferritinophagy- and mitochondrial-related proteins in H9C2 cells following JNK inhibition under ISO and PP2A inhibition conditions.
