## Supplemental Material for "Protein Phosphatase 2A Activation Attenuates Acute Myocardial Injury in Takotsubo Syndrome by Modulating Ferroptosis and Mitochondrial Injury in Cardiomyocytes"

**Methods**

**Bulk RNA-seq analysis from public datasets**

To investigate conserved stress-responsive signaling pathways associated with Takotsubo syndrome (TTS), publicly available bulk RNA-seq datasets were obtained from the Gene Expression Omnibus (GEO) database. A rat stress cardiomyopathy RNA-seq dataset (GSE223385) generated from ventricular tissues subjected to immobilization stress was analyzed to identify transcriptional programs associated with stress-induced cardiac injury. In addition, transcriptomic data generated from a transverse aortic constriction (TAC)-induced stress cardiomyopathy model in potassium channel Kv1.5-deficient mice were analyzed to assess conserved signaling alterations in stress-induced cardiac dysfunction. Differentially expressed genes between the control and stress cardiomyopathy groups were identified and subsequently subjected to Gene Ontology (GO) enrichment analysis to determine biological pathways associated with catecholamine-induced myocardial stress. For pathway-level visualization and comparison, significantly enriched biological processes were ranked by enrichment scores and adjusted P values.

**Proteomic analysis of public human dataset**

To determine whether stress-responsive signaling alterations identified in experimental models were conserved in human TTS, the publicly available human proteomic dataset (GSE95368) from patients with stress cardiomyopathy was analyzed. Differentially expressed proteins between control and TTS samples were subjected to pathway enrichment analysis to identify biological processes associated with stress-induced cardiac injury. Pathway enrichment and data visualization were performed using R software.

**scRNA-seq analysis of public dataset**

To characterize cellular heterogeneity and stress-responsive signaling alterations in TTS at single-cell resolution, publicly available single-cell RNA sequencing (scRNA-seq) datasets were obtained from the GEO database under accession number GSE305273. This dataset included cardiac cells isolated from control and isoproterenol (ISO)-treated mouse hearts. Raw count matrices were processed and analyzed using the R package Seurat. This processing encompassed steps such as quality control, dimensionality reduction, and clustering. Cell clusters were annotated based on the expression of established lineage-specific marker genes. Comparative analyses between control and ISO-treated hearts were performed to evaluate alterations in cellular distribution and relative cell-type composition under catecholamine stress conditions. Differentially expressed genes between control and ISO-treated groups, as well as among specific cell populations, were identified using the Seurat package. GO enrichment analyses were subsequently performed to determine stress-responsive biological pathways associated with catecholamine-induced cardiac injury. In addition, expression patterns of PP2A-related genes, including Ppp2ca, were analyzed across distinct cardiac cell populations. The proportion of Ppp2ca-positive cells and the relative expression levels within individual clusters were quantified and compared between experimental groups. Statistical significance for differential expression analysis was determined using false discovery rate (FDR)-adjusted P values, with an adjusted P<0.05 considered statistically significant.

**Human plasma samples**

Human plasma samples from patients with TTS were obtained at Emory University Hospital under protocols approved by the Institutional Review Board of Emory University. Plasma samples from individuals without known cardiovascular or other diseases served as controls. A total of 5 control and 5 TTS plasma samples were included in this study. Blood samples were collected in EDTA-containing tubes, centrifuged to isolate plasma, aliquoted, and stored at -80°C until use.

**Animals**

All protocols involving animals were approved by an Institutional Animal Review Committee at Emory University and were performed in compliance with the Guide for the Care and Use of Laboratory Animals. Male C57BL/6J mice aged 16 weeks were purchased from the Jackson Laboratory (Bar Harbor, ME). Cardiac-specific PP2A-Cα deficient mice were generated by crossing the floxed Ppp2cα mice with Myh6-Cre^+/-^ mice (The Jackson Laboratory, strain no. 005657). Cardiac-specific Ppp2cα deficient was achieved by administering 2 consecutive doses of tamoxifen (50μl, 20 mg/mL, dissolved in sunflower seed oil; Sigma-Aldrich) through intraperitoneal injection to 14-week-old male Ppp2cα flox^+/-^-Myh6-Cre^+/-^ mice (CM-PP2Afl^+/-^). Age-matched male Ppp2cα-floxed mice without Cre expression and Myh6-Cre^+/-^ mice, subjected to the same tamoxifen regimen, were used as controls. Following a 2-week washout period, 16-week-old CM-PP2Afl^+/-^ mice were used for the modeling of TTS.

**Animal Model of TTS**

To induce TTS in mice, 16-week-old male mice were intraperitoneally (i.p.) injected with ISO (400 mg/kg, Sigma-Aldrich) for 2 hours, 7 days, and 14 days, as described previously.^30,41^ To further validate and enhance the reliability of the findings, a second TTS model was established using 10-week-old male C57BL/6J mice, which received a single intraperitoneal injection of EPI (2.5 mg/kg, Sigma-Aldrich) for 2 hours as previously described.^33^ SMAPs (5mg/kg of DT-061) were administered via oral gavage 10 minutes after acute ISO or EPI treatment, whereas the control mice received the same vehicle solution. A working solution for DT-061 was prepared in an N,N-dimethylacetamide (DMA)/Kolliphor HS-15 (Solutol)/ddH_2_O solution. The PP2A inhibitor LB-100 (1.0 mg/kg, Selleck Chemicals) was administered through i.p. injection 24h prior to acute ISO treatment. To further validate the PP2A-mediated signaling pathway involved in TTS, mice were pretreated with LB-100 (2 mg/kg, i.p.) and/or the JNK inhibitor SP600125 (5 mg/kg, MedChem Express) dissolved in DMSO, 24 hours before ISO administration. Follow-up echocardiography was performed, and the mice were humanely euthanized using CO_2_ asphyxiation at designated time points according to their assigned groups. The hearts were then immediately harvested for further analysis.

**Echocardiography**

Transthoracic echocardiography was performed on anesthetized mice using a Vevo 3100 ultrasound imaging system (FUJIFILM Visual Sonics, Canada) with an MX550D transducer. An optimal parasternal long-axis view was acquired to trace morphological changes in the left ventricular (LV) apical myocardium. All parameters of systolic function were obtained from the short-axis under M-mode tracings at the level of the papillary muscle. A cut-off of over 500 beats per minute was used to avoid the effects of narcosis on cardiac function, and three consecutive short-axis cardiac cycle parameters were used for calculation.

**RNA sequencing and data analysis**

Total RNA was extracted from the apex of hearts from Vehicle-treated or DT-061-treated control and TTS mice (4 groups, n=4 per group) using Trizol and RNeasy mini kits (Qiagen). All samples that passed RNA quality control were subjected to commercial RNA-Seq (Novogene Corporation Inc.) using the NovaSeq PE150 platform, with a sequencing depth of 6 G of raw data per sample. Differentially expressed genes (DEGs) were defined as |log2 (fold change)| > 1.5 and p value < 0.05. Heatmaps of DEGs were generated using Heatmap package (version 1.0.8) in R (<https://cran.r-project.org/web/packages/pheatmap/index.html>). Gene set enrichment analysis for KEGG and GO was performed using Metascape (<http://metascape.org/gp/index.html#/main/step1>) to identify the biological pathways and processes affected in the apex of hearts associated with DT-061 treatment in TTS mice.

**Immunofluorescence staining**

For the in vivo samples, paraffin sections were permeabilized with 0.3% Triton X-100 for 10 minutes after antigen retrieval and blocked with 3% BSA for 30 minutes at room temperature. For the in vitro samples, H9C2 cells and primary human cardiac myocytes were seeded on cell slides in 24-well plates, received the corresponding treatment, then fixed with 4% paraformaldehyde, and subsequently permeabilized and blocked with 3% BSA. Next, the above samples were incubated overnight at 4°C with primary antibodies α-Actinin (Cell Signaling, 13522S) 1:50, mL309 (7C10) 1:50, and 4-HNE (Thermo Scientific, MA5-27570) 1:500. Sections and slides were then incubated with host-specific secondary antibodies with Alexa Fluor 488 or 594 for one hour. Finally, DAPI was used to counterstain the nuclei. Images were captured at a resolution of 1024 × 1024 pixels and 8 bits using a Zeiss LSM 800 Airyscan Laser Scanning Confocal Microscope (Zeiss, Oberkochen, Germany) and quantified using ImageJ software. Sections or cell slides incubated solely with fluorescently conjugated secondary antibodies served as negative controls.

**Histology and immunohistochemistry (IHC)**

Mice were euthanized, and the hearts were perfused with ice-cold PBS via the left ventricle. The heart tissues were harvested, rinsed in PBS, sliced into apical, mid-ventricular, and basal sections, fixed in 4% paraformaldehyde for 48 h, and embedded in paraffin. Serial sections (5 μm) from each group were stained with hematoxylin and eosin (H&E) for morphological analysis. For immunohistochemical staining, dewaxed and hydrated sections, following antigen retrieval, were incubated with 3% hydrogen peroxide for 15 minutes and then blocked with 5% goat serum for one hour at room temperature. Next, Sections were incubated overnight with mL309 (7C10) (1:50) at 4°C under humidified conditions for three nights, followed by incubation for one hour with secondary anti-mouse immunoglobulins/HRP (1:200, Vector Laboratories) at room temperature. Finally, the sections were stained with a DAB kit and then counterstained with hematoxylin. Control slides stained with species-matched IgG instead of the primary antibody were prepared to determine antibody specificity. Images were captured using a NanoZoomer (Hamamatsu) and processed with NDP.veiw 2.

**Transmission electron microscopy**

Samples from the apex (1 mm ×2 mm ×2 mm) were quickly removed from the left ventricle and immediately fixed in 2.5% glutaraldehyde in 0.1M cacodylate buffer (pH 7.4) for one day at room temperature. The Emory University Robert P. Apkarian Integrated Electron Microscopy Core Facility then post-fixed, embedded, sectioned, and mounted the samples. The samples were then imaged using a JEOL 1400EX electron microscope at 80 kV (JEOL). At least five fields of view were randomly selected for each sample.

**Cell culture**

Primary human cardiac myocytes (HCM) (PromoCell, C-12810), derived from the left ventricles of the adult heart, were cultured in myocyte growth medium (PromoCell, C-22070). Cardiomyocytes at passage 5–8 were used for subsequent studies. H9C2 cells (ATCC, CRL-1446), derived from embryonic rat myocardium, were cultured in high-glucose DMEM supplemented with 10% fetal bovine serum (FBS). To establish an in vitro catecholamine stress model of TTS, HCM and H9C2 cells were serum-starved overnight and subsequently stimulated with ISO for 2 hours. For dose-response experiments, H9C2 cells were treated with ISO at concentrations of 1, 5, or 10 mM for 2 hours.^30^ Based on preliminary experiments demonstrating robust suppression of methyl-PP2A-C at 10 mM ISO, this concentration was used in subsequent mechanistic studies unless otherwise indicated. For pharmacological modulation of PP2A activity, cells were pretreated with the PP2A activator DT-061 (5 μM) for 6 hours or with the PP2A inhibitor LB-100 (2 μM) for 2 hours before ISO stimulation. For autophagic flux inhibition experiments, cells were pretreated with bafilomycin A1 (BafA1; 100 nM) for 1 hour before ISO treatment. For JNK inhibition studies, cells were co-treated with LB-100 and the JNK inhibitor SP600125 (10 μM) for 2 hours before ISO exposure. To assess the effects of circulating stress-associated factors from patients with TTS, human plasma samples were diluted in baseline culture medium and applied to cardiomyocytes for 2 hours to assess methyl-PP2A-C expression.

**siRNA-mediated gene silencing**

For the small interfering RNA (siRNA)-mediated knockdown study, H9C2 cells were cultured in 6-well plates and transfected with siRNAs targeting NCOA4, FTMT, or with control siRNA (Horizon Discovery) using Lipofectamine RNAiMAX reagent (Invitrogen) at a concentration of 30 nM. Eight hours after transfection, the medium was replaced, and cells were cultured for an additional 60 hours before harvesting. Total protein was extracted to assess siRNA transfection efficiency by Western blot analysis.

**Cell death staining**

H9C2 cardiomyocytes were seeded in Glass Bottom Culture Dishes (Matsunami Glass, D35-14-1.5-U, USA). The following day, cells were transfected with siRNA and incubated for 60 hours. Subsequently, cells were pre-treated with DT-061 (5 μM) or LB-100 (2 μM), followed by treatment with or without ISO for 2 hours. In some experiments, cells were co-treated with SP600125 (10 μM) and LB-100. Cardiomyocyte death was assessed using the Live/Dead Viability/Cytotoxicity Assay Kit (Proteintech) at 37°C for 30 minutes. Images were acquired on a Zeiss LSM 800 Airyscan Laser Scanning Confocal Microscope (Zeiss, Oberkochen, Germany).

**Detection of lipid peroxidation**

H9C2 cardiomyocytes were seeded into the 35 mm glass bottom dishes. Treated cells were washed three times with sterile Hank's Balanced Salt Solution (HBSS) and then incubated with 10 μM BODIPY™581/591 C11 in serum-free DMEM for 30 minutes at 37°C following the manufacturer's instructions. Dishes were then washed, and fluorescence was imaged on a Zeiss LSM 800 Airyscan Laser Scanning Confocal Microscope (Zeiss, Oberkochen, Germany).

**Detection of intracellular and mitochondrial Fe^2+^**

To assess cellular Fe*^2+^*levels, H9C2 cardiomyocytes were plated on glass bottom culture dishes and treated as described. Cardiomyocytes were then washed with HBSS and incubated with 1 µM FerroOrange (Dojindo) in serum-free medium for 30 minutes at 37°C in 5% CO₂. Fluorescence images were immediately acquired on a Zeiss Confocal Microscope (Zeiss, Oberkochen, Germany). For mitochondrial iron detection, H9C2 cells and primary human cardiac myocytes were washed three times with HBSS and incubated with 5 μM Mito-FerroGreen and MitoTracker Red (Dojindo) for 30 minutes at 37°C, following the manufacturer's instructions. Cells were then washed three times with HBSS and examined using a confocal microscope. Relative fluorescence intensity was analyzed using ImageJ.

**Mitochondrial lipid peroxide detection**

Mitochondrial lipid peroxides were detected using the cell-permeable fluorescent probe MitoPeDPP (3-[4-(perylenylphenylphosphino)phenoxy]propyltriphenylphosphonium iodide) (Dojindo). Cells were inoculated in a glass-bottom dish and incubated with MitoPeDPP working solution (0.5 μM) for 15 min. The solution was removed, and the cells were washed twice with PBS. The cultures were then pretreated with DT-061 for 6 hours or LB-100 for 2 hours, followed by ISO stimulation. After rinsing with PBS, the cells were examined using a confocal fluorescence microscope.

**Detection of mitochondrial membrane potential**

H9C2 cardiomyocytes and primary human cardiac myocytes were seeded in 24-well plates. After treatment, the culture medium was replaced with a JC-1 working solution (Dojindo) and cells were incubated at 37°C for 30 minutes. Cells were then washed twice with HBSS, and imaging buffer solution was added. Fluorescence images were captured using a microscope at excitation wavelengths of 488 nm (green) and 594 nm (red) and quantified using ImageJ software.

**Evaluation of mitochondrial morphology**

H9C2 cells were seeded in the 35 mm glass bottom dishes. Treated cells were gently washed three times with HBSS and incubated with 0.1 μM MitoTracker Deep Red (Dojindo) in serum-free DMEM for 30 minutes at 37°C. Cells were washed and subsequently examined using a Zeiss LSM 800 Airyscan Laser Scanning Confocal Microscope (Zeiss, Oberkochen, Germany). Fluorescence intensity was quantified using ImageJ.

**Western blot analysis**

For total protein extraction, myocardial tissues from the apex or cells were lysed in 1x RIPA lysis buffer (Invitrogen, Carlsbad, CA, USA) supplemented with Halt protease and phosphatase inhibitor cocktail (Thermo Fisher Scientific) and centrifuged at 12,000g (15 minutes, 4°C). Supernatants were collected, and protein concentrations were measured using the Pierce BCA Protein Assay Kit (Thermo Fisher Scientific, 23227) according to the manufacturer’s instructions. Total proteins or cytoplasmic and mitochondrial proteins were separated on 4%-12% SDS-PAGE gels and subsequently transferred to PVDF membranes (Millipore, Billerica, MA, USA). The PVDF membranes were blocked with 5% BSA in Tris-buffered saline containing 0.05% Tween-20 (TBST), then incubated overnight at 4°C with primary antibodies, followed by horseradish peroxidase-conjugated goat anti-rabbit or anti-mouse secondary antibody. Antibodies and dilutions: PTGS2 (Cell Signaling, 12282S) 1:1000, SLC7A11 (Cell Signaling, 12691S) 1:1000, GPX4 (Cell Signaling, 59735S) 1:1000, FTH1 (Cell Signaling, 4393S) 1:500, FTMT (Invitrogen, MA568234) 1:500, P62 (Cell Signaling, 39749S) 1:1000, LC3B (Cell Signaling, 2775S) 1:500, Pgc1α (ABclonal, A12348) 1:1000, TOM20 (Proteintech, 66777-1-Ig) 1:1000, TIM23 (Proteintech, 11123-1-AP), NLRP3 (Proteintech, 68102-1-Ig) 1:500, Bax (Cell Signaling, 2772S) 1:1000, Bcl-2 (Proteintech, 68103-1-Ig) 1:1000, GSDMD-N (Cell Signaling, 10137S) 1:1000, GSDMD (Proteintech, 20770-1-AP) 1:1000, C-caspase-3 (Cell Signaling, 9661S) 1:1000, Caspase-3 (Cell Signaling, 9662S) 1:1000, P-JNK (Cell Signaling, 4668P) 1:1000, JNK (Cell Signaling, 9252S) 1:1000, P-P38 (Cell Signaling, 4511S) 1:1000, P38 (Cell Signaling, 9212S) 1:1000, Erk (Cell Signaling, 4696S) 1:1000, P-Erk (Cell Signaling, 4370S) 1:1000, Vinculin (Proteintech, 66305-1-Ig) 1:10000, HRP anti-rabbit IgG goat secondary antibody (Cell Signaling, 7074S), and HRP anti-mouse IgG goat secondary antibody (Cell Signaling, 7076S). Target proteins were visualized using Immobilon Western HRP substrate (Millipore Sigma) on the Bio-Rad ChemiDoc Western Blot Imager. Total proteins were normalized to Vinculin. The quantitative analysis of protein bands was performed using the Image J software.

**Statistical Analysis**

Statistical analyses were performed using GraphPad Prism version 9.0 (GraphPad Software Inc, San Diego, CA, USA). Data from the quantitative experiments are presented as mean ± standard error from the mean (SEM) of at least 3 independent experiments. For comparisons between 2 groups, an unpaired 2-tailed Student's t test was performed for normally distributed data, and the Mann-Whitney test for non-normally distributed data. For comparisons among multiple groups, one-way ANOVA (followed by Tukey post hoc test) was applied. Survival rates were assessed using Kaplan-Meier analysis. *P* < 0.05 was considered statistically significant.
